# Deubiquitinating enzyme mutagenesis screens identify a USP43 driven HIF-1 transcriptional response

**DOI:** 10.1101/2024.01.10.574971

**Authors:** Tekle Pauzaite, Niek Wit, Rachel V Seear, James A Nathan

**Affiliations:** Cambridge Institute of Therapeutic Immunology & Infectious Disease (CITIID), Jeffrey Cheah, Biomedical Centre, Department of Medicine, University of Cambridge, Cambridge, CB2 0AW

**Keywords:** Hypoxia, USP43, HIF, deubiquitination, DUB, oxygen-sensing, 14-3-3, deubiquitylation

## Abstract

The ubiquitination and proteasome-mediated degradation of Hypoxia Inducible Factors (HIFs) is central to metazoan oxygen-sensing, but the involvement of deubiquitinating enzymes (DUBs) in HIF signalling is less clear. Here, using a bespoke DUBs sgRNA library we conduct CRISPR/Cas9 mutagenesis screens to determine how DUBs are involved in HIF signalling. Alongside defining DUBs involved in HIF activation or suppression, we identify USP43 as a DUB required for efficient activation of a HIF response. USP43 is hypoxia regulated and selectively associates with the HIF-1α isoform, and while USP43 does not alter HIF-1α stability, it facilitates HIF-1 nuclear accumulation and binding to its target genes. Mechanistically, USP43 associates with 14-3-3 proteins in a hypoxia and phosphorylation dependent manner to increase the nuclear pool of HIF-1. Together, our results unveil the DUB landscape in HIF signalling, and highlight the multifunctionality of DUBs, illustrating that they can provide important signalling functions alongside their catalytic roles.

## Introduction

The ability of organisms to sense and adapt to varying oxygen gradients is conserved across species. In metazoans, oxygen-sensing is principally governed by HIFs, which facilitate an adaptive transcriptional programme to respond to reduced oxygen availability (Ivan & Kaelin, 2017; Kaelin & Ratcliffe, 2008; Schofield & Ratcliffe, 2004; Semenza, 2012). The oxygen-sensitive nature of the HIF pathway relates to prolyl hydroxylation of the HIF-α subunit (principally HIF-1α or HIF-2α). When oxygen is abundant, the HIF-α subunit undergoes prolyl hydroxylation at two conserved residues by the prolyl hydroxylases (PHDs or EGLNs), which prime HIF-α for ubiquitination by the Von Hippel– Lindau (VHL) E3 ligase, leading to its rapid proteasome-mediated degradation (Bruick & McKnight, 2001; Epstein *et al*, 2001; Jaakkola *et al*, 2001; Maxwell *et al*, 1999). When oxygen supply is limited PHDs are inactivated and HIF-α is stabilised, thereby allowing the formation of the stable HIF-α/HIF1β heterodimeric transcriptional complex which binds to hypoxia responsive elements (HREs) at HIF responsive genes.

The dominance of VHL-mediated ubiquitination in controlling HIF-α stability is evident from extensive genetic studies of *VHL* mutations, and compounds inhibiting VHL enzymatic activity (Buckley *et al*, 2012; Cancer Genome Atlas Research, 2013; Frost *et al*, 2016; Latif *et al*, 1993; Maher & Kaelin, 1997; Maxwell *et al*., 1999; Mitchell *et al*, 2018; Turajlic *et al*, 2018). In all cases, VHL loss or inhibition leads to HIF-α stabilisation, even when oxygen levels are abundant. VHL-independent ubiquitination has been also observed (Ferreira *et al*, 2015; Flügel *et al*, 2012), and this may help regulate HIF-α levels when oxygen supply is limited, or to fine tune the HIF response. However, while it is evident that ubiquitination of HIF-α is required for oxygen-sensing, the role of deubiquitination and deubiquitinating enzymes (DUBs) is less clear.

HIF-α deubiquitination has been proposed as a potential regulatory mechanism to counteract VHL, with several DUBs linked to reversing HIF-α stabilisation, including USP7, USP8, USP20, USP29 and USCHL1 (Gao *et al*, 2021; Li *et al*, 2005; Troilo *et al*, 2014; Wu *et al*, 2016). Generally, HIF-α deubiquitination does not overcome the dominant action of VHL when oxygen is present, and is mostly observed by overexpression of the DUB or re-oxygenation experiments, or in VHL-deficient kidney cancer cells (Hong *et al*, 2020). However, it is important to consider that DUBs may be involved in other aspects of HIF signalling, and not just HIF-α stability (Bett *et al*, 2013; Bremm *et al*, 2014; Moniz *et al*, 2015). Moreover, DUBs may have additional roles outside of their catalytic activity (Nishiya *et al*, 2011), which may be equally important in regulatory pathways.

Here, we take an unbiased approach to comprehensively explore the contribution of DUBs to HIF signalling. Using CRISPR/Cas9 mutagenesis screens and a dynamic fluorescent reporter that provides a robust readout of HIF signalling, we conduct both suppressor and activator screens to define the principal DUBs involved in HIF regulation. Alongside defining the global involvement of DUBs involved in HIF regulation, our screening approach identified USP43, a poorly characterised DUB that was required for activation of the HIF-transcriptional response. RNA-seq corroborated the involvement of USP43 in HIF signalling, with USP43 as one of only two DUBs that is upregulated in hypoxia in a HIF-dependent manner. Functionally, USP43 depletion decreases the activation of HIF-1 target genes, without altering HIF-2 signalling. This selectivity of USP43 for HIF-1 was not due to altered HIF-α protein or mRNA levels. Instead, we show that USP43 associates with HIF-1α and promotes the nuclear accumulation of HIF-1 and the chromatin binding of the transcriptional complex to specific HIF-1 target genes. Remarkably, USP43 facilitates HIF-1 nuclear accumulation through a hypoxia-sensitive recruitment of 14-3-3 proteins, rather than its deubiquitinase activity. Therefore, this work demonstrates the ability of a DUB to influence a transcriptional programme and highlights the broader role of DUBs in intracellular signalling pathways.

## Results

### Mutagenesis screens identify USP43 as regulator of the HIF response

To unbiasedly identify deubiquitinating enzymes (DUBs) involved in HIF regulation we used a phenotypic HIF functional screen using a pooled DUB CRISPR sgRNA library. The screens were performed in HeLa cells expressing a well validated HIF reporter (HIF^ODD^GFP reporter), which we have previously used to interrogate regulation of the HIF response (Bailey *et al*, 2020; Burr *et al*, 2016; Ortmann *et al*, 2021). This reporter has the advantage that it provides a dynamic readout to both endogenous HIF stability and transcriptional activation via binding to a triplicate HRE (**Fig 1A**). The sub-pooled DUB sgRNA library was generated as part of a ‘ubiquitome library’, encoding 10 sgRNAs per gene (Menzies *et al*, 2018). We performed two opposing mutagenesis screens, aimed at identifying DUBs that either activate or suppress the HIF response (**Fig 1A**) (**Supplemental Data 1**). OTUD5 was significantly enriched in the suppressor screen (**Fig 1B**), whereas both USP43 and USP52 (also known as PAN2) were enriched in the activator screen (**Fig 1C**). USP52 has been implicated in HIF regulation, through its global effect on RNA stability, and we validated this finding (**Fig S1A**) (Bett *et al*., 2013; Wolf & Passmore, 2014), confirming our approach. OTUD5 (also known as DUBA) has been implicated in multiple cell pathways, including mTOR regulation (Cho *et al*, 2021), and OTUD5 depletion only had a marginal change in HIF^ODD^GFP reporter levels in 21 % oxygen (**Fig S1B**). However, the involvement of USP43, a poorly characterised DUB, in the HIF response was unknown, and we confirmed that USP43 depletion prevented full activation of the HIF^ODD^GFP reporter in 1 % oxygen (**Fig 1D, E; S1A**), validating the screen findings.

**Fig 1.**
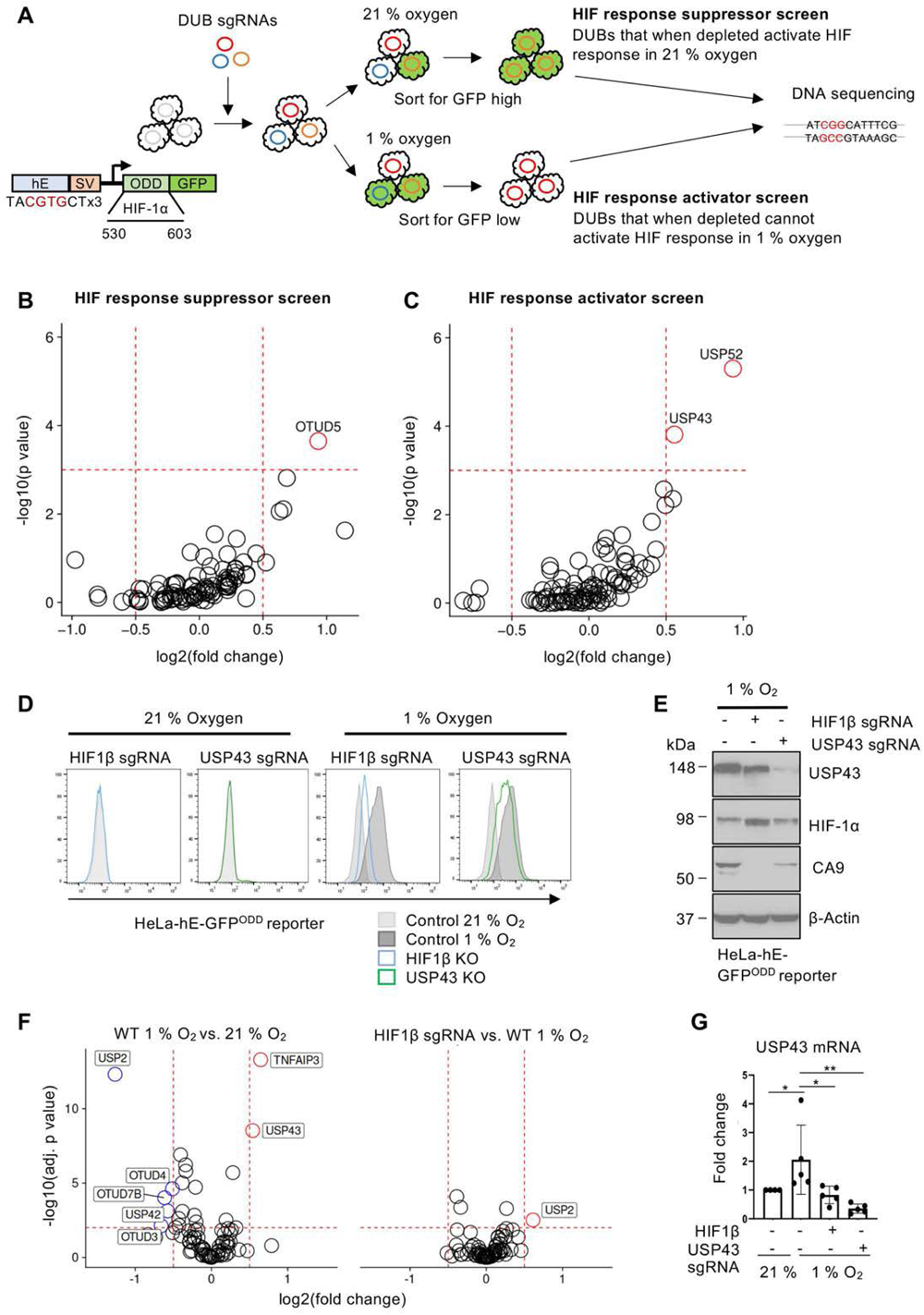
Mutagenesis screens identify USP43 as regulator of the HIF response. **A)** Schematic of the pooled DUB sgRNA library screen. Diagram of HIF-HRE-GFP^ODD^ reporter in HeLa-Cas9 cells (left). For the suppressor screen, HeLa HRE-^ODD^GFP reporter cells were transduced with a DUB sgRNA library, and sorted for a GFP high population using flow cytometry. For the activator screen, cells were incubated in 1 % oxygen for 24 h and sorted for GFP low expressing cells. **B, C)** Comparative bubble plots of HIF suppressor screen **(B)**, or activator screen **(C)**. DNA was extracted, sgRNAs identified by Illumina MiniSeq, and compared between sorted and unsorted libraries. Unadjusted P value calculated using MaGECK robust rank aggregation (RRA). **D)** HeLa HRE-GFP^ODD^ reporter expression in mixed knockout populations (denoted as sgRNA) of HIF1β (light blue) or USP43 (green). Reporter levels were analysed by flow cytometry for GFP 488 nm following incubation in 21 % or 1 % oxygen (16 h). **E)** Immunoblot of HIF1β or USP43 depleted HeLa HRE-GFP^ODD^ reporter cells after 16 h of 1 % oxygen. Representative of three biological replicates. **F)** RNA-seq analysis of HeLa control or a mixed KO population of HIF1β KO cells, treated with either 21 % or 1 % oxygen for 16 h. n= 3 biologically independent samples. Volcano plots of differential mRNA expression in control 1 % versus 21 % oxygen (left), or HIF1β KO cells compared with control cells in 1 % oxygen (right). A log2(fold change) of > 0.5 or < -0.5, and – log10(p value) of > 2.5 was selected as the significance level. **G)** Quantitative RT-PCR (qPCR) of USP43 mRNA in HeLa cells. Mixed KO population of HIF1β or USP43 HeLa cells were incubated in 1 % or 21 % oxygen for 16 h. n=5 biologically independent samples. Mean ± sd, *P≤ 0.05, **P≤ 0.01, two-way ANOVA.

We reasoned that DUBs that are regulatory in HIF signalling may be reciprocally regulated by HIFs, as observed by other components of HIF pathway (Minamishima *et al*, 2009; Ortmann *et al*., 2021). Therefore, using RNA-seq (**Fig S1C, Supplementary Data 2**), we analysed which DUBs are transcriptionally regulated in hypoxia, and whether this is dependent on HIFs. HeLa cells exposed to 1 % or 21 % oxygen for 16 hours demonstrated that few DUBs had altered expression in hypoxia (**Fig 1F**). *OTUD7B* (also known as Cezanne) was the top downregulated DUB in hypoxia, consistent with its known involvement in non-canonical HIF stability (Bremm *et al*., 2014; Moniz *et al*., 2015). *USP43* was transcriptionally upregulated in hypoxia, and this hypoxic induction of *USP43* was not observed in HIF1β null cells (**Fig 1F, G**), indicating that *USP43* expression was HIF inducible. As USP43 was both a top hit in the screens and was itself regulated in a HIF-dependent manner, we focused our further studies on the actions of this DUB.

### USP43 is required for activation of a HIF-1 response

We first determined if USP43 was required for endogenous HIF signalling, and not just the HIF^ODD^GFP reporter. Mixed knockout (KO) populations of USP43 using sgRNA in multiple cell lines (HeLa, A549, MCF7) showed reduced protein levels of Carbonic Anhydrase 9 (CA9) – a well validated HIF-1 target (**Fig 2A**). The reduction in CA9 was not to the same extent as completely ablating the HIF pathway by HIF1β loss, but CA9 levels were reduced without any appreciable difference in protein levels of the two main HIF-α isoforms (HIF-1α, HIF-2α) or HIF1β (**Fig 2A; S2A, B**). USP43 clonal loss both delayed the HIF driven CA9 induction and decreased the amplitude of the HIF response (**Fig 2B, C; S2C-F**). Similar findings were observed using dimethyloxalylglycine (DMOG), a broad-spectrum inhibitor of PHDs and 2-oxoglutarate dependent dioxygenases (**Fig 2D; S2B**), indicating that USP43 was likely involved in HIF signalling downstream of prolyl hydroxylation. Moreover, qPCR analysis in A549, HeLa, and MCF-7 cells confirmed that USP43 depletion decreased the expression of selected HIF target genes but not expression of HIF-1α or HIF-2α (**Fig 2E; S3A-C**).

**Fig 2.**
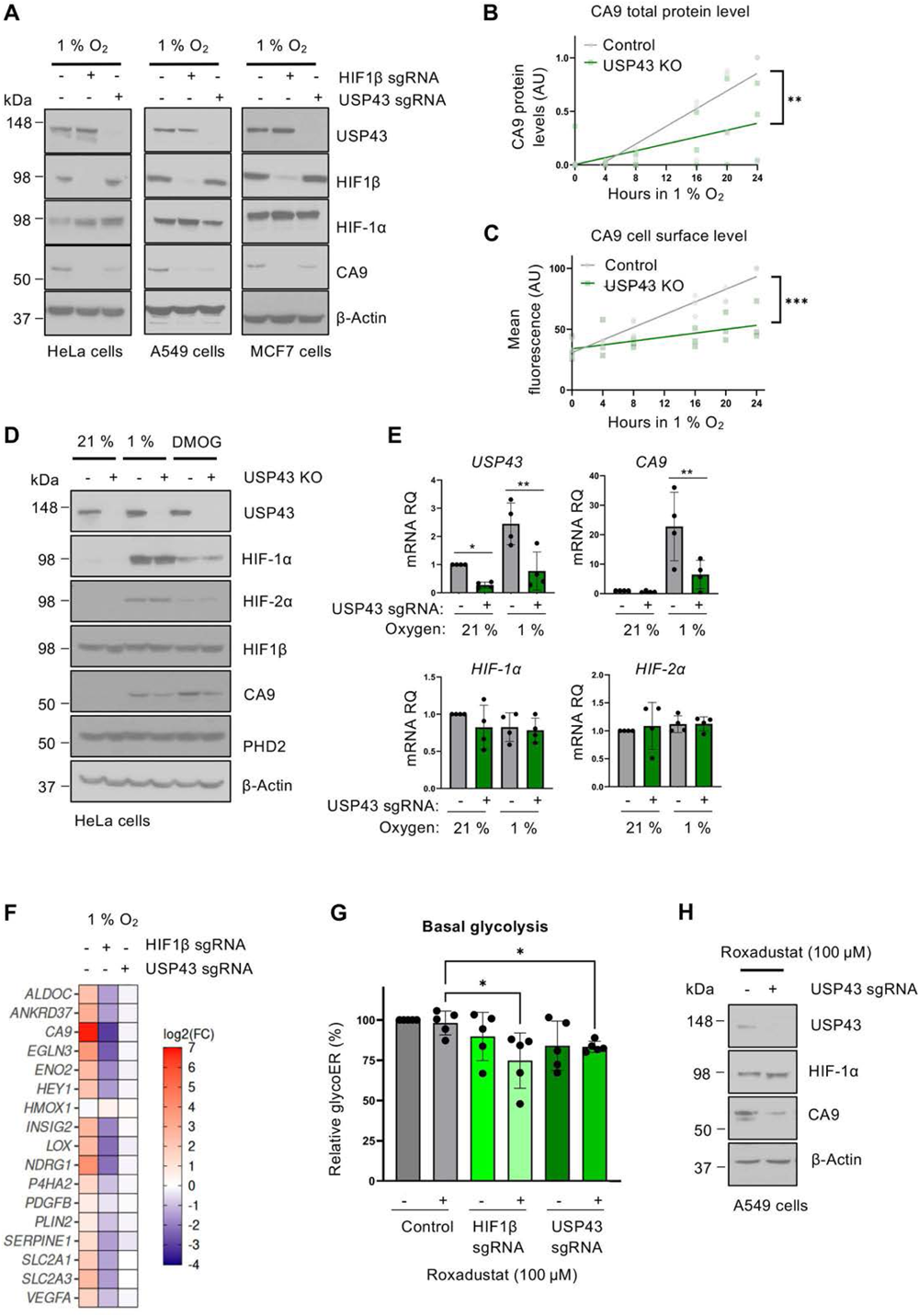
USP43 is required for activation of a HIF-1 response. **A)** USP43 depletion in different cell types. Mixed KO populations of USP43 or HIF1β HeLa, A549, MCF7 cells were generated. Cells were incubated in 1 % oxygen for 16 hr and immunoblotted. Representative of three biological replicates. **B, C)** Quantification of total **(B)** and cell surface **(C)** CA9 protein levels in control or USP43 null (KO denotes null clone) HeLa cells after 0-24 h of 1 % oxygen. Total protein levels measured by immunoblot densitometry relative to a β-actin control (n=3 biological replicates, **P≤ 0.01, two-way ANOVA). Cell surface CA9 levels measured by flow cytometry, and the mean fluorescence intensity is shown (n=3 biological replicates, ***P≤ 0.001, two-way ANOVA). Representative immunoblots and flow cytometry shown in **Fig S2C-F. D)** Immunoblot of HeLa-Cas9 or USP43 null clone after 16 h of 1 % oxygen or 24 h of 1 mM DMOG treatment. Representative of three biological replicates. **E)** qPCR of *USP43*, *CA9*, *HIF-1α*, or *HIF-2α* in control or USP43 depleted HeLa cells following incubation in 1 % or 21 % oxygen for 16 h (n=4 biologically independent samples, mean ± sd, *P≤ 0.05, **P≤ 0.01, two-way ANOVA). **F)** Heat map representing mRNA expression of validated HIF target genes analysed by RNA sequencing in control, HIF1β or USP43 mixed KO HeLa cells. **G, H)** Bioenergetic analysis. Control, HIF1β depleted or USP43 depleted A549 cells were incubated with or without Roxadustat (0.1 mM) for 24 h. Cells were analysed using a Seahorse XF analyser by performing a Glycolytic Rate Assay **(G)**. The protein efflux rate derived from glycolysis (glycoPER) as a percentage of total were calculated using WAVE version 2.6.1. (n=5, *P≤ 0.05, two-way ANOVA). Immunoblot of samples used for the used for Seahorse analysis **(H)**. Control or USP43 depleted A549 cells, incubated with or without Roxadustat (0.1 mM) for 24 h.

To substantiate the involvement of USP43 in HIF-transcriptional activation of target genes, we undertook RNA-seq analysis in HeLa control, USP43 depleted, or HIF1β depleted cells following 6 hr of incubation in 21 % or 1 % oxygen (**Fig 2G; S1C**). Principal component analysis (PCA) showed that all experimental conditions clustered similarly in 21 % oxygen, indicating that USP43 or HIF1β loss did not globally alter transcription when oxygen was present (**Fig S1C**). USP43 depletion also clustered with the control HeLa cells exposed to 1 % oxygen, rather than the HIF1β deficient cells where the HIF transcriptional response is completely ablated. These findings suggested that USP43 loss either had a moderate effect on HIF target gene expression or altered a subset of HIF target genes. Indeed, the expression analysis of HIF target genes showed a partial reduction in transcript levels when compared with HIF1β KO in 1 % oxygen, and interestingly, USP43 loss altered the transcription of genes involved in glycolysis (**Fig 2F**). Bioenergetic analysis using a Seahorse confirmed that USP43 loss decreased basal glycolysis following PHD inhibition with Roxadustat (**Fig 2G, H**). Therefore, USP43 loss perturbs the HIF response in a selective manner, and can impair the HIF-driven shift to glycolysis.

Given that USP43 only regulated a subset of HIF-target genes, we investigated whether USP43 had any specificity towards HIF-α isoforms. Using HeLa HIF^ODD^GFP reporter cells that were either HIF-1α or HIF-2α deficient (Ortmann *et al*., 2021), we observed that USP43 loss only altered HIF-1 transcriptional activation of the reporter (**Fig 3A, B**). These findings were substantiated using HIF-1α or HIF-2α null HeLa cells, whereby we observed a reduction in the activation of HIF-1 but not HIF-2 target genes (**Fig S4A-D**). To further validate these findings, we examined the effect of USP43 loss in clear cell renal carcinoma cell (ccRCC) lines that have constitutive activation of both HIF-1 and HIF-2 (RCC4 cells, *HIF-1α*+, *HIF-2α*+, and *VHL*-), or only encode HIF-2α (786-0 cells, *HIF-1α*-, *HIF-2α*+, and *VHL*-). USP43 depletion reduced CA9 protein, transcript and cell surface levels in RCC4 cells (**Fig 3C-E**), but did not influence HIF-2α driven transcription in 786-0 cells (**Fig 3F, G**). Therefore, USP43 shows specificity towards the HIF-1α isoform.

**Fig 3.**
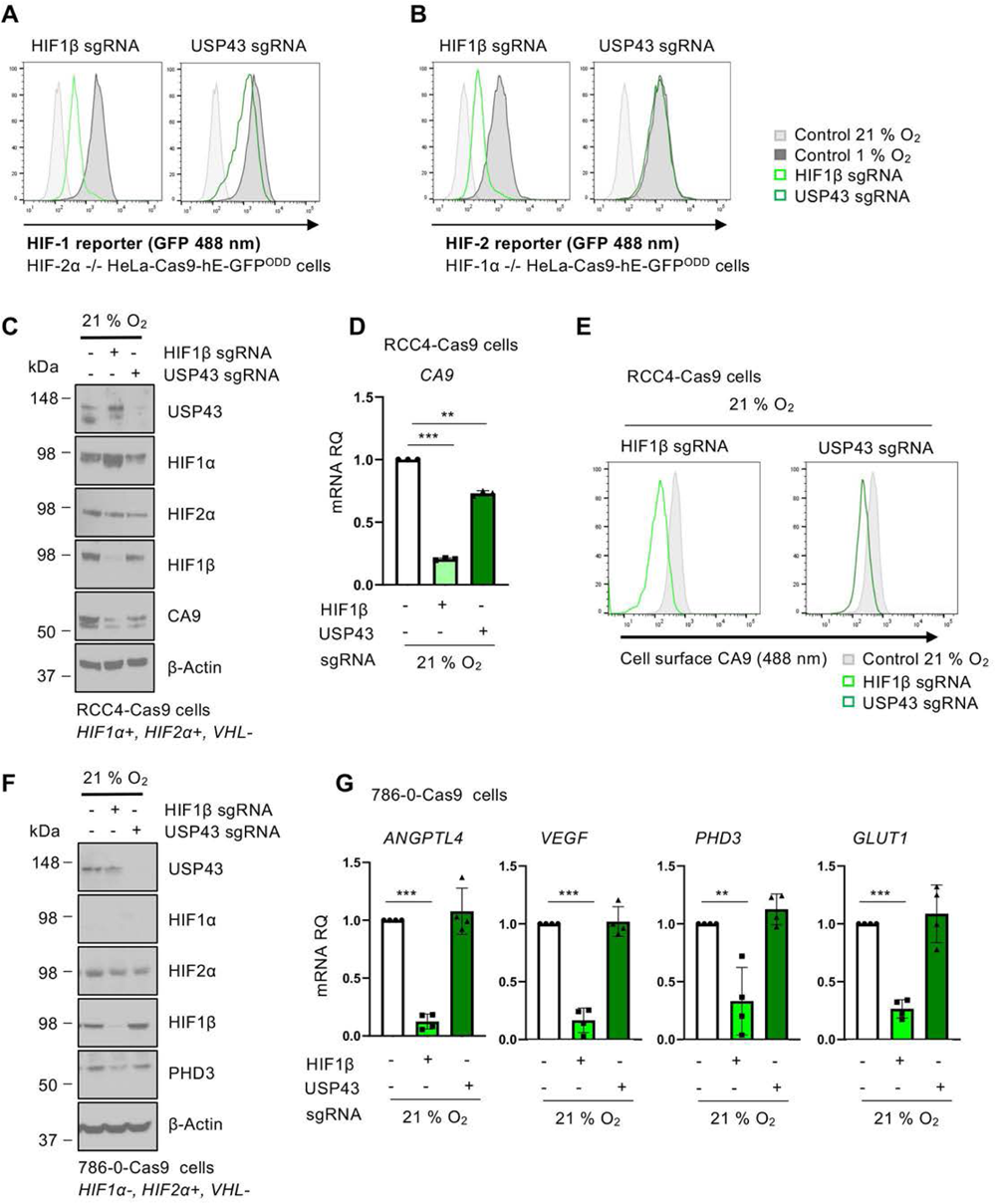
USP43 is specific for the HIF-1 complex. **A, B)** Mixed KO populations of either USP43 or HIF1β in HIF-1 reporter cells (HIF-2α null HeLa HRE-^ODD^GFP) **(A)** or HIF-2 reporter cells (HIF-1α null HeLa HRE-^ODD^GFP) **(B)** were generated, treated with 1 % or 21 % oxygen for 24 h, and reporter activity measured by flow cytometry. Representative of three biological replicates. **C-E)** USP43 depletion in RCC4 cells. Mixed KO populations of USP43 or HIF1β were generated in RCC4 cells stably expressing Cas9. HIF complex component levels and CA9 levels were measured by immunoblot **(C).** *CA9* expression **(D)** and CA9 cell surface protein levels **(E)** were analysed by qPCR and flow cytometry respectively. Representative of three biological replicates. Mean ± sd, *P≤ 0.05, ***P≤ 0.001, two-way ANOVA. **F, G)** USP43 depletion in 786-0 cells. USP43 mixed KO populations were generated in 786-0 cells stably expressing Cas9. HIF complex components and HIF-2 target genes were analysed by immunoblot **(F)** or qPCR **(G)** as previously described. n=4 biological replicates. Mean ± sd, **P≤ 0.01, ***P≤ 0.001, two-way ANOVA.

### USP43 is an active DUB that associates with the HIF-1 complex

To understand how USP43 may facilitate a HIF-1 transcriptional response, we examined if USP43 associated with HIF-1α in hypoxia by incubating HeLa cells in 21 % or 1 % oxygen for 6 hr and immunoprecipitating endogenous USP43 or HIF-1α. USP43 bound to HIF-1α with either the USP43 pull-down or the HIF-1α pull-down, but not to HIF-2α (**Fig 4A, B; S5A**)

**Fig 4.**
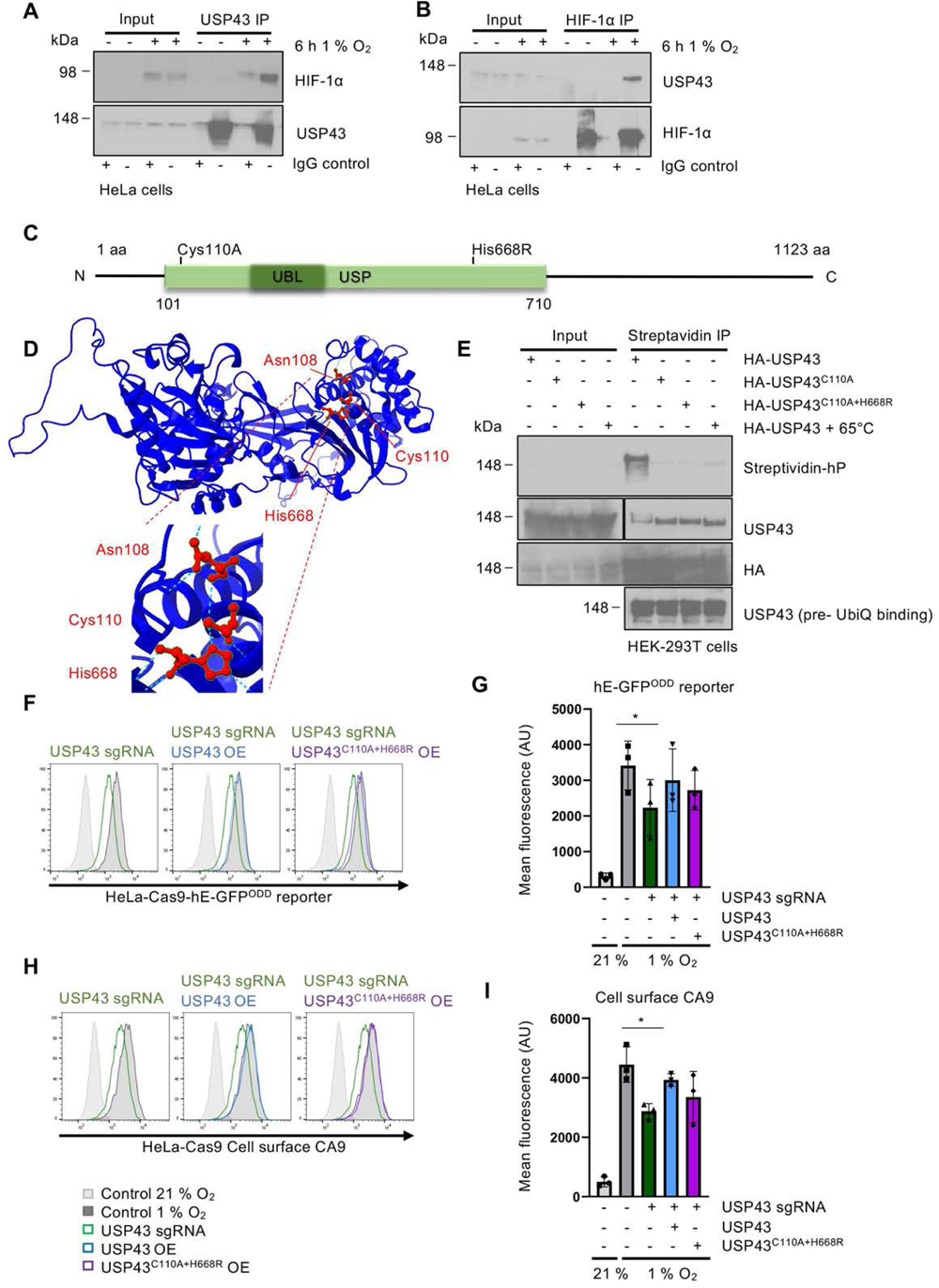
USP43 is an active DUB that associates with the HIF-1 complex. **A)** Endogenous USP43 was immunoprecipitated in HeLa cells grown at 21 % or 1 % oxygen for 6 h. Samples were immunoblotted for HIF-1α. Rabbit IgG was used a control for non-specific binding. Representative of 3 biological replicates. **B)** As in **(A)** but endogenous HIF-1α was immunoprecipitated and samples were immunoblotted for endogenous USP43. **C)** Schematic of USP43 protein sequence, with ubiquitin specific protease (USP) and ubiquitin-like domains (UBL) shown. The positions of the active site nucleophile cysteine 110 (Cys110) and proton acceptor histidine 668 (His668) are indicated. **D)** Predicted USP43 catalytic triad using AlphaFold (10.1093/nar/gkab1061) and ChimeraX-1.6.1.exe. The catalytic site Cys110, His668, and Asn108 are labelled. **E)** Immunoblot of DUB catalytic activity-based probe Biotin-ANP-Ub-PA (UbiQ-077) (1 µM) binding to the overexpressed HA-USP43, HA-USP43^C110A^, or HA-USP43^C110A+H668R^ in HEK-293T cells. **F, G)** HeLa HRE-^ODD^GFP reporter cells or USP43 mixed population KO cells (dark green) reconstituted with HA-USP43 (blue) or HA-USP43^C110A+H668R^ (purple) after 16 h of 1 % oxygen treatment. HIF reporter activity measured by flow cytometry **(F)** and quantified **(G)** using FlowJo10. Each dot represents the mean fluorescence intensity (AU at 488 nm) of a biological replicate (n=3, *P≤ 0.05, two-way ANOVA). **H, I)** As for **(F, G)** but for CA9 cell surface levels in HeLa cells (n=3, *P≤ 0.05, two-way ANOVA).

As USP43 associated with HIF-1α it was possible that it directly regulated HIF-1 complex ubiquitination. However, it was unlikely that USP43 controlled HIF-1α stability, as HIF-1α protein levels were not altered at steady state (**Fig 2A, D**), and we confirmed that neither USP43 depletion nor overexpression altered HIF-1α degradation kinetics using cycloheximide chase assay of HIF-1α degradation (**Fig S5B-E**).

Given the lack of effect of USP43 on HIF-1α stability and that the deubiquitinase activity of USP43 has not been well studied, we considered whether USP43 was an active DUB or a pseudoDUB as observed for other USP proteins (Bett *et al*., 2013; Wolf & Passmore, 2014). We generated mutations of the putative catalytic triad of USP43 (nucleophilic cysteine 110 and proton acceptor histidine 668) based on the Alphafold predictions (Jumper *et al*, 2021) (**Fig 4C, D**), and used a biotinylated DUB probe (Biotin-ANP-Ub-PA) that covalently binds the active cysteine to investigate DUB activity. Wildtype USP43 was able to bind the ubiquitin probe, demonstrating that USP43 is catalytically active (**Fig 4E**), but mutation of cysteine 110 to alanine (C110A), with or without a histidine 668 to arginine (H668R) mutation rendered USP43 catalytically inactive.

To determine the involvement of USP43 deubiquitinase activity in HIF signalling, we tested if reconstitution of the catalytically active or inactive USP43 restored HIF^ODD^GFP reporter or CA9 levels following USP43 depletion. USP43 expression restored HIF activity in HeLa and A549 USP43 deficient cells (CRISPR/Cas9 KO or siRNA) in a dose dependent manner, similarly to control conditions (**Fig 4F-I; S4F-I**). Surprisingly, the USP43^C110A+H667R^ mutant also partially restored HIF reporter activity and CA9 levels (**Fig 4F-I**), suggesting that while USP43 enzymatic activity may be involved, deubiquitination itself was not essential for the effect of USP43 on HIF-mediated transcription.

### USP43 regulates HIF-1α nuclear accumulation

We postulated that USP43 must act downstream of HIF-1α stabilisation, and was involved in either subcellular localisation of HIF-1, or the interaction of the transcription factor with chromatin. We therefore examined where USP43 was localised within the cell, and if this altered under hypoxic conditions. Immunofluorescence of endogenous USP43 was not possible, but using HA-USP43 we observed that USP43 was predominantly localised within the cytosol under conditions of both 21 % and 1 % oxygen (**Fig 5A**). However, some USP43 localised within the nucleus, and this was markedly increased in hypoxia (**Fig 5A, B**). We confirmed that endogenous USP43 also localised to the nucleus using biochemical fractionation of HeLa cell lysates (**Fig 5C; S6A, B**). This translocation of USP43 resulted in a rapid increase in USP43 within the chromatin fraction and corresponded with the chromatin-associated HIF-1α, and still occurred when the HIF-1 complex was absent (**Fig 5C; S6A, B**). Using a more stringent isolation of chromatin, involving formaldehyde cross-linking and urea extraction (Kustatscher *et al*, 2014), we still observed USP43 in the chromatin fraction, and noticed that HIF-1α enrichment in the chromatin fraction was reduced in USP43 KO cells and increased when USP43 was overexpressed (**Fig 5D, F**). Therefore, we examined if USP43 altered the kinetics of HIF-1α localisation to the nucleus in hypoxia. USP43 deficient HeLa cells, incubated in 1 % oxygen for 20, 40 and 60 min showed reduced HIF-1α levels on chromatin, whilst USP43 overexpression increased HIF-1α levels in the chromatin fraction (**Fig 5E, G; S6C, D**). Overexpression of the USP43 catalytic inactive mutant (USP43^C110A+H667R^) did not increase HIF-1α in the chromatin fraction (**Fig S6E, F**). Therefore, while both catalytic and non-catalytic activity of USP43 may be involved, our data are consistent with a role for USP43 in HIF nuclear localisation or retention.

**Fig 5.**
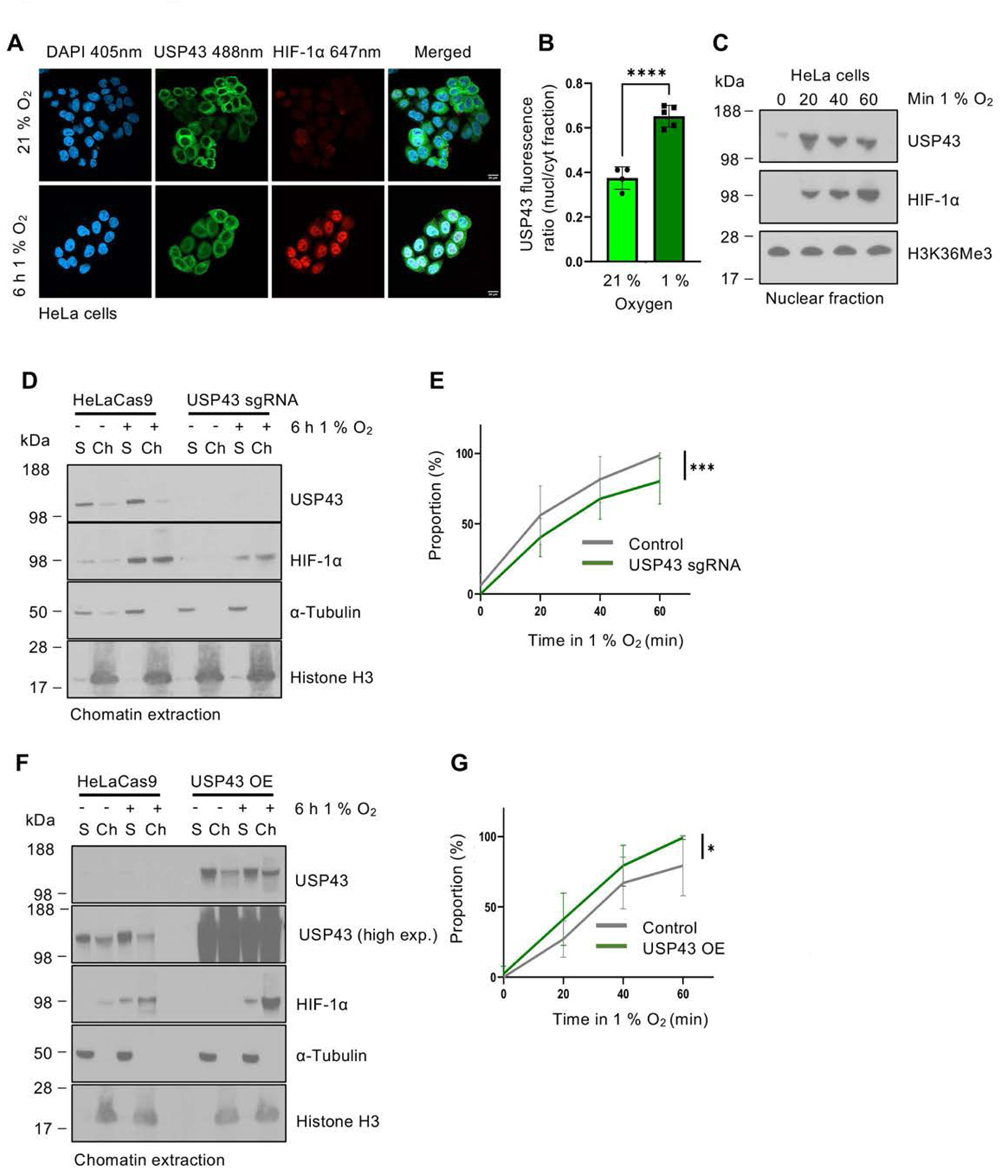
USP43 regulates HIF-1α nuclear accumulation. **A, B)** Confocal immunofluorescence microscopy of USP43 overexpressing HeLa cells stained for USP43 (488 nm) and HIF-1α (647 nm). Cells were grown at 21 % or 1 % oxygen for 6 h on coverslips prior to fixation. Representative images **(A)** and quantification **(B)** shown. Each point represents a biological replicate of approximately 25 cells acquired and analysed as one image. Analysis was performed blinded on ImageJ-win64. n=5 biologically independent samples, mean ± sd, ****P≤ 0.0001, two-way ANOVA. **C)** Immunoblot of the nuclear fraction of HeLa-Cas9 incubated in 1 % oxygen for 0 to 60 min. **D)** Immunoblot of the soluble (S) and chromatin fractions (Ch) obtained using a formaldehyde cross-linking chromatin extraction protocol. HeLa-Cas9 and USP43 mixed population KO cells were incubated in 21 % or 1 % oxygen for 6 h prior to lysis. **E)** Quantification of HIF-1α enrichment within the nuclear fraction in HeLa control or USP43 mixed population KO cells. Cells were incubated in 1 % oxygen for 0 to 60 min prior to lysis, and HIF-1α levels relative to a stable histone mark (H3K36me3) were measured by immunoblot. Representative immunoblot shown in **Fig S6C**. n=3 biologically independent samples, mean ± sd, ***P≤ 0.001, two-way ANOVA. **F)** As for **(D)** but with HeLa control and USP43 overexpression (OE). Representative immunoblot shown in **Fig S6D**. **G)** As for **(E)** but with HeLa control and USP43 overexpression. n=3 biologically independent samples, mean ± sd, *P≤ 0.05, two-way ANOVA.

Given that USP43 levels perturbed the kinetics of the abundance of HIF-1α on chromatin, we next determined if this altered the binding of HIF-1α to known target gene loci in HeLa cells. Chromatin immunoprecipitation (ChIP)-PCR of HIF-1α binding showed that USP43 depletion decreased HIF-1α binding to HREs on the *CA9 and PHD3* loci in hypoxia (**Fig 6A**). The decrease in HIF-1α binding was not as substantial as HIF1β depletion, consistent with the functional consequences of USP43 loss described previously. USP43 loss also did not alter HIF-1α binding to the *VEGF* HRE, similarly to the lack of effect of USP43 depletion on *VEGF* using RNA-seq. Comparable findings were observed using HIF1β ChIP-PCR (**Fig 6B**), indicating that USP43 depletion altered binding of the HIF-1 complex at selected HIF-1 loci. Interestingly, these findings were distinct from other known co-activators involved in the regulation of HIF target genes, such as the SET1B histone methyltransferase, as we observed no changes in binding of the HIF-1 complex to chromatin with SET1B loss, but both USP43 and SET1B loss still resulted in reduced histone 3 lysine 4 tri-methylation (H3K4me3) at the *CA9* locus (**Fig 6C, D**). Moreover, we did not observe any changes in Histone 2B lysine120 ubiquitination (H2BK120Ub) following USP43 loss, nor an association between USP43 and H2BK120Ub, in contrast to a prior report implicating USP43 in the regulation of this activating epigenetic mark (He *et al*, 2018) (**Fig S6G, H**).

**Fig 6.**
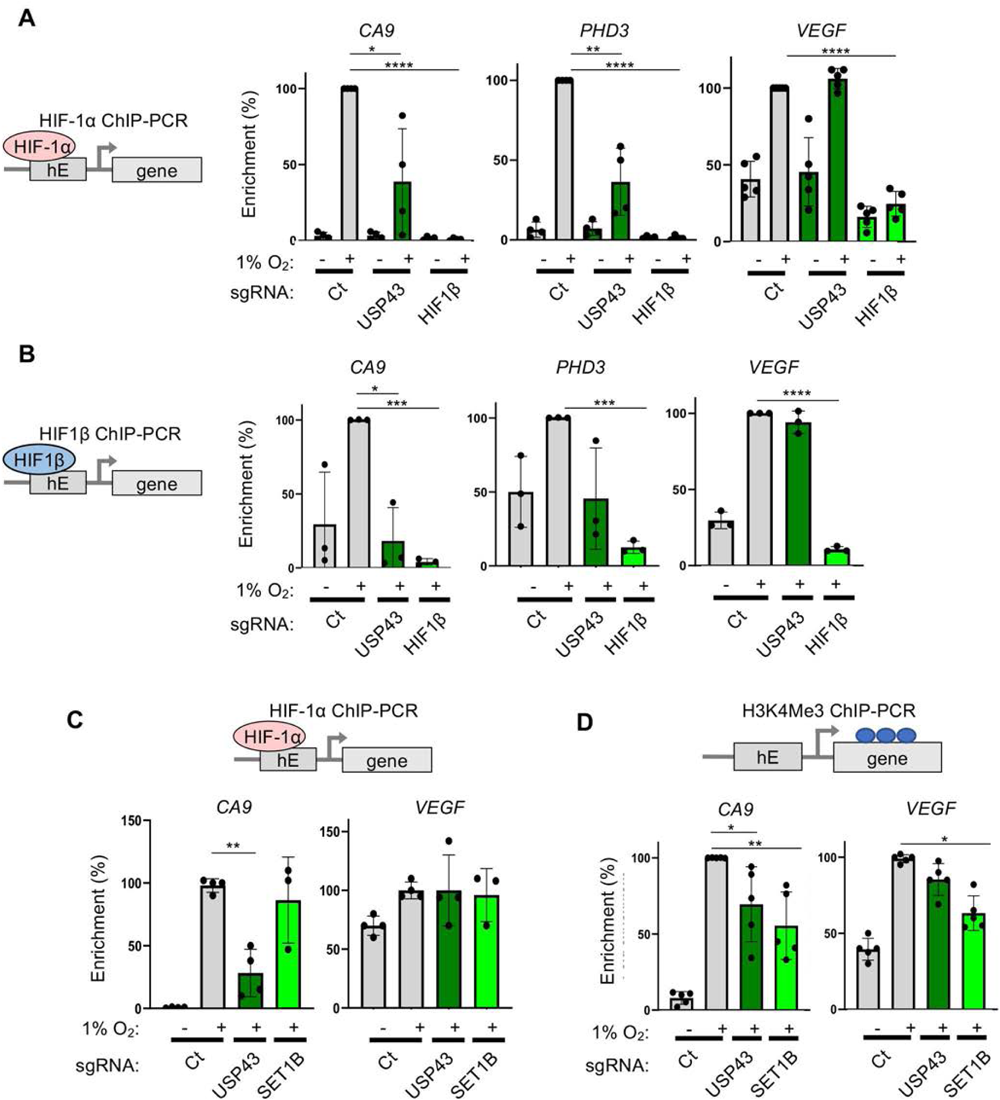
USP43 KO reduces HIF-1 binding to chromatin at HREs. **A)** HIF-1α ChIP-PCR for HREs of *CA9, PHD3*, and *VEGF* in control, HIF1β or USP43 depleted HeLa cells treated with 21 % or 1 % oxygen for 6 h. n=4 biologically independent samples, mean ± sd, *P≤ 0.05, **P≤ 0.01, ****P≤0.0001, two-way ANOVA). **B)** as **(A)** but HIF1β binding to specific HREs. n=3 biologically independent samples, mean ± sd, *P≤ 0.05, ***P≤ 0.001, ****P≤0.0001, two-way ANOVA. **C)** HIF-1α ChIP-PCR at *CA9* or *VEGF* HREs in HeLa control, USP43 depleted (n=4 biological repeats) or SET1B depleted (n=3 biological repeats) HeLa cells. Cells were incubated in 21 % or 1 % oxygen for 6 h prior to lysis. **D)** As for **(C)** but immunoprecipitated for H3K4me3 at the HREs of *CA9* or *VEGF* (n=4 biological repeats). mean ± sd, *P≤ 0.05, **P≤ 0.01, two-way ANOVA.

### USP43 associates with 14-3-3 proteins to regulate HIF-1 signalling

The ability of USP43 levels to influence HIF-1α levels on chromatin suggested that a further protein or signal was involved. We therefore used the global DUB mass spectrometry interactome (Sowa *et al*, 2009) to find potential USP43 binding partners. USP43 was the only DUB shown to strongly interact with 14-3-3 proteins (Sowa *et al*., 2009), and given the importance of 14-3-3 in nuclear-cytoplasmic transport (Brunet *et al*, 2002) and potential HIF-1 interactions (Peng et al., 2022; He et al., 2018; Tang et al., 2016; Xue at al., 2016), we hypothesised that USP43 may regulate HIF-1 localisation through the recruitment of 14-3-3 proteins. We confirmed that USP43 binds 14-3-3 proteins, using overexpressed HA-USP43 and endogenous USP43 in HeLa cells (**Fig 7A, B**), consistent with the DUB mass spectrometry interactome (Sowa *et al*., 2009). Moreover, we found that 14-3-3 proteins bound USP43 more strongly in 1 % oxygen (**Fig 7B, C**).

**Fig 7.**
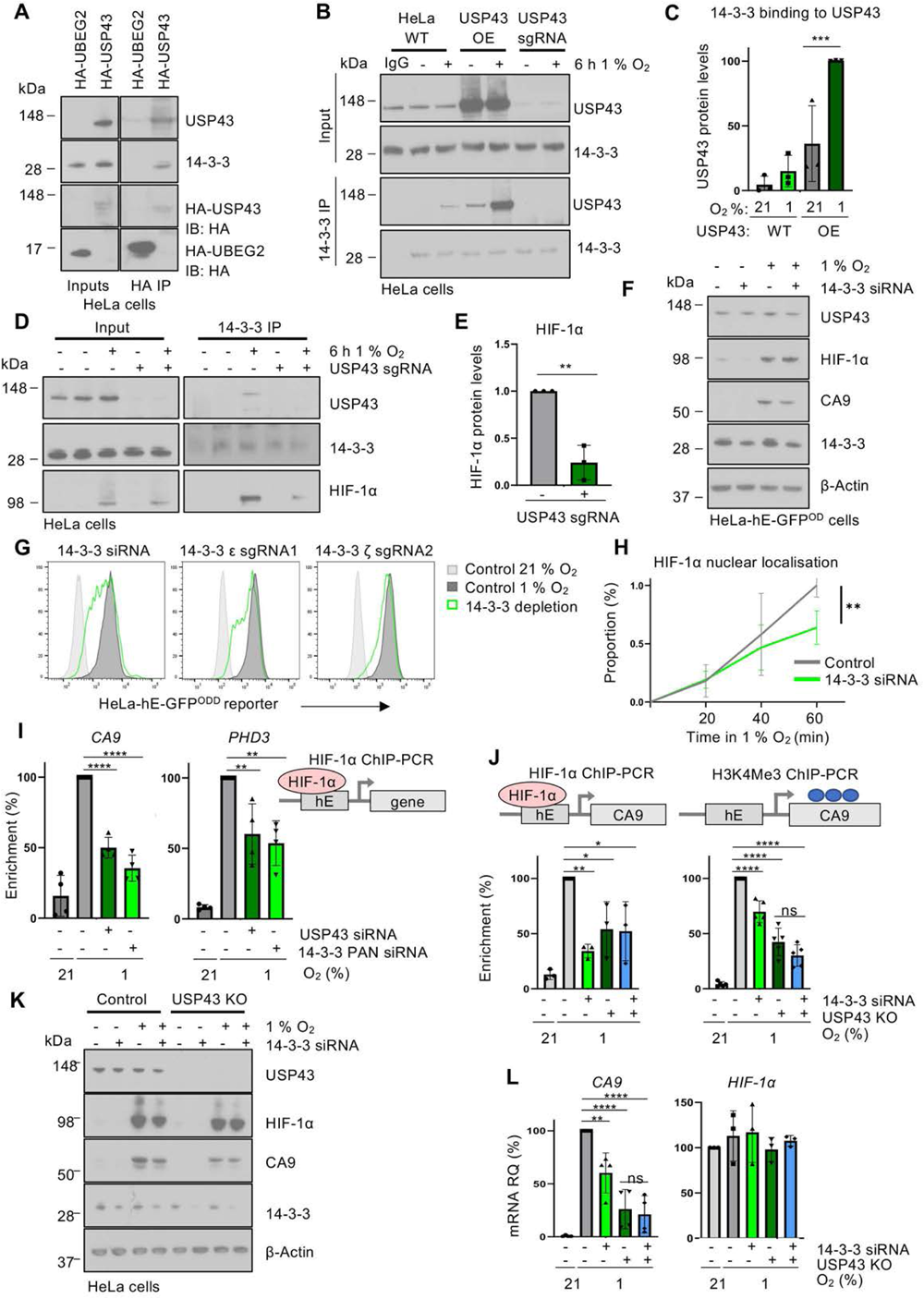
USP43 associates with 14-3-3 to regulate HIF-1 signalling. **A)** HeLa cells stably expressing HA-USP43 or HA-UBEG2 were immunoprecipitated and immunoblotted using a pan 14-3-3 antibody (recognises all isoforms). **B, C)** Control, USP43 depleted (sgRNA) or overexpressing HeLa cells were incubated at 21 % or 1 % oxygen for 6 h prior to lysis. Endogenous 14-3-3 proteins were immunoprecipitated and samples were immunoblotted for USP43 **(B)**. Quantification of USP43 associated with immunoprecipitated 14-3-3 **(C)**. USP43 protein levels normalised to USP43 input. n=3 biological replicates, mean ± sd, ***P≤ 0.001, two-way ANOVA. **D, E)** Control or USP43 depleted HeLa cells were incubated in 21 % or 1 % oxygen for 6 h prior to lysis. Endogenous 14-3-3 was immunoprecipitated and immunoblotted for USP43 and HIF-1α **(D)**. Quantification of HIF-1α association with 14-3-3 proteins (relative to HIF-1α input protein level) **(E)**. n=3 biological replicates, mean ± sd, **P≤ 0.01, two-way ANOVA. **F)** Immunoblot of HeLa HRE-^ODD^GFP reporter and 14-3-3 siRNA treated cells after 16 h of 21 % or 1 % oxygen treatment. Immunoblot representative of three biological replicates. **G)** HeLa HRE-^ODD^GFP reporter cells treated with 14-3-3 pan siRNA, 14-3-3 epsilon sgRNA or 14-3-3 zeta sgRNA were analysed by flow cytometry. Control levels of the reporter in 21 % (light grey) or 1 % (dark grey) oxygen are shown. **H)** Quantification of HIF-1α enrichment within the nuclear fraction in HeLa control or following 14-3-3 siRNA-mediated depletion. Cells were incubated in 1 % oxygen for 0 to 60 min prior to lysis, and HIF-1α levels relative to a stable histone mark (H3K36me3) were measured by immunoblot. Representative immunoblot shown in **Fig S7J**. n=3 biologically independent samples, mean ± sd, **P≤ 0.01, two-way ANOVA. **I)** HIF-1α ChIP-PCR for HREs of *CA9 or PHD3* in control, USP43 or 14-3-3-siRNA depleted HeLa cells treated with 21 % or 1 % oxygen for 6 h. n=4 biologically independent samples, mean ± sd, **P≤ 0.01, ****P≤0.0001, two-way ANOVA). **J-L)** Control or USP43 null HeLa cells, with or without 14-3-3 depletion, were incubated in 21 % or 1 % oxygen for 16 h. Samples were analysed by ChIP-PCR for HIF-1α (n=3) or H3K4me3 (n=5) at the HRE of *CA9* **(J)**, immunoblot of HIF-1α, 14-3-3, CA9 and USP43 (representative of 3 biological replicates) **(K)** or qPCR for *CA9* (n=4) and *HIF-1α* (n=3) mRNA expression **(L)**. Mean ± sd, *P≤ 0.05, **P≤ 0.01, ****P≤0.0001, two-way ANOVA.

14-3-3 proteins are well known to interact with phosphorylated proteins. Consistently, we showed that USP43 is phosphorylated using Lambda protein phosphatase (Lambda PP) and a gel shift assay (**Fig S7A**). We also observed that immunoprecipitation with a phospho-Serine (pSer) 14-3-3 motif antibody pulled down USP43, and importantly, this interaction was hypoxia dependent (**Fig S7B**). Moreover, the interaction between 14-3-3 and USP43 could be replicated using Culyculin A as a phosphatase inhibitor (Resjö *et al*, 1999), and conversely, prevented by Lambda PP (**Fig S7C**).

We next examined if the USP43 and 14-3-3 interaction was important for the recruitment of HIF-1α, and observed that USP43 loss decreased the levels of HIF-1α associated to endogenous 14-3-3 (**Fig 7D, E**). Additionally, using a pan-siRNA mediated depletion of 14-3-3 (targeting 6 of 7 isoforms) (**Fig S7D**), we observed reduced HIF activation of the fluorescent HIF reporter and endogenous CA9, without affecting HIF-1α protein and mRNA levels (**Fig 7F, G; S7E, F**). These data supported the hypothesis that the USP43 association with 14-3-3 was important for regulating HIF-1 signalling.

To substantiate the involvement of 14-3-3 proteins, we attempted to map the 14-3-3 interaction site but with 168 potential binding sites identified (Madeira *et al*, 2015), further investigation of the specific 14-3-3 binding sites by site directed mutagenesis was unfeasible. We therefore focused on which 14-3-3 isoform was involved. We generated sgRNA targeting all seven 14-3-3 isoforms and observed the largest effect on HIF-1 signalling following depletion of zeta (14-3-3ζ) and epsilon (14-3-3ε) (**Fig 7G; S7G**). These findings were consistent with the anti-14-3-3 pan siRNA predominantly targeting 14-3-3ζ and 14-3-3ε (**Fig 7G; S7F**). Furthermore, when we overexpressed HA-14-3-3ζ or 14-3-3ε, we found HA-14-3-3ζ bound more USP43 in 1 % oxygen compared to 21 % oxygen (**Fig S7H, I**).

If the binding of USP43 to 14-3-3 isoforms was important for HIF-1 signalling, we would expect a defect in HIF-1α binding to chromatin and that 14-3-3 and USP43 function within the same pathway. In support of this, we observed that pan siRNA-mediated depletion of 14-3-3 proteins reduced the levels of HIF-1α by chromatin fractionation (**Fig 7H; S7J**) and decreased the binding of HIF-1α to the HREs of *CA9* and *PHD3*, similarly to USP43 depletion (**Fig 7I**). Lastly, we confirmed that both USP43 and 14-3-3 function in the same pathway by using siRNA-mediated depletion of 14-3-3 in HeLa USP43 null clones. 14-3-3 depletion did not further reduce HIF-1α binding to the *CA9* locus or H3K4me3 levels in the USP43 KO cells (**Fig 7J**). Moreover, *CA9* mRNA expression and protein levels were reduced the to the same extent in the USP43 null clones irrespective of combined depletion of 14-3-3 (**Fig 7K, L; S7K**). Together, these data confirm that USP43 helps mediate the interaction between HIF-1α and 14-3-3ζ/ε, which in turn supports the chromatin localisation of HIF-1α and activation of HIF-1 target genes (**Fig 8**).

**Fig 8.**
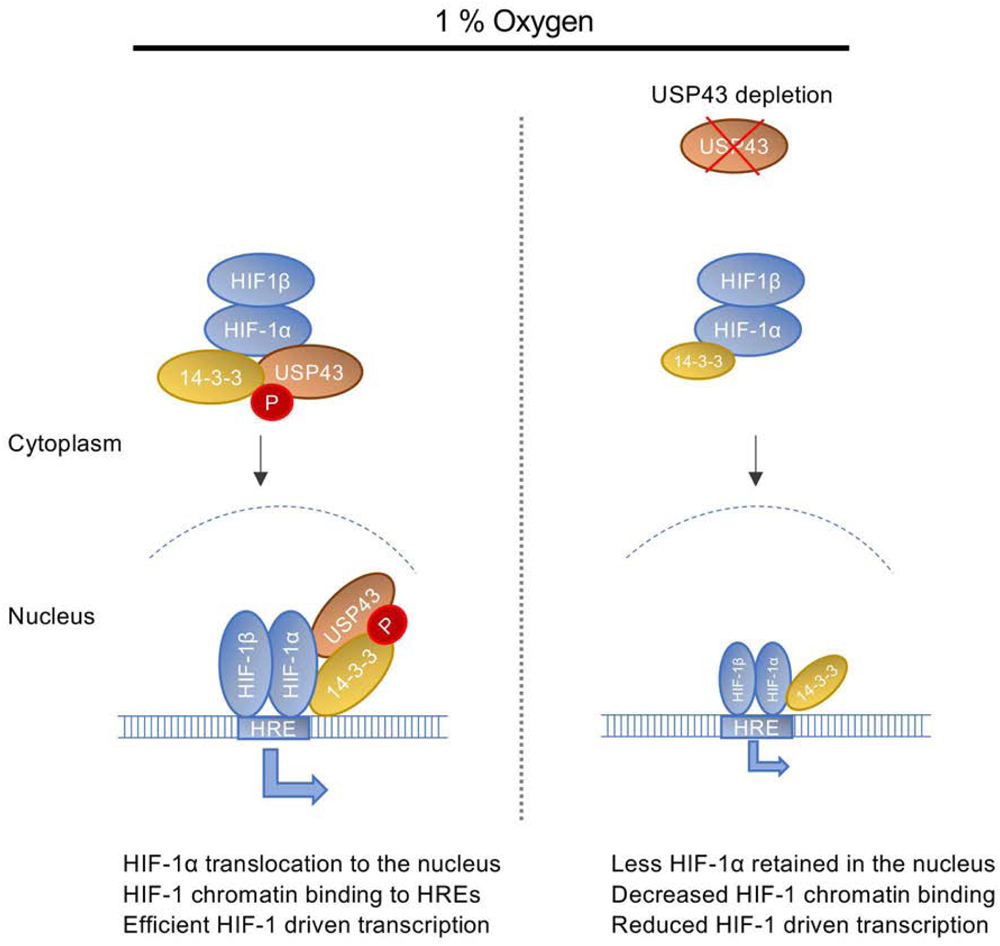
USP43 mediated regulation of a HIF-1 transcriptional response. Schematic model of the role of USP43 in the HIF response. In hypoxia, USP43 associates with the stabilised HIF-1α in a phosphorylation and 14-3-3 dependent manner. This USP43 association promotes HIF-1 accumulation in the nucleus, and is required for the efficient of HIF-1 to HREs on chromatin to activate gene transcription.

## Discussion

Our approach of using a pooled DUB sgRNA library allowed us to unbiasedly screen for DUBs involved in the HIF response. This strategy proved powerful in identifying USP43 and its involvement in HIF-1 activation through associating with 14-3-3ζ/ε. It may seem surprising that USP43 catalytic activity was not absolutely required, but our findings highlight that DUBs may also serve important non-catalytic roles.

USP43 is the only DUB found to strongly associate with 14-3-3 proteins in an unbiased mass spectrometry interactome (Sowa *et al*., 2009), and given the multifunctional roles of 14-3-3 proteins, this interaction likely helps facilitate USP43 function, or localise USP43 within the cell. The predominant effect of USP43 on the HIF-1 transcriptional response was through association with 14-3-3ζ/ε. USP43 has been reported to associate with other 14-3-3 isoforms, including a 14-3-3β/ε heterdimer (He *et al*., 2018). It is therefore plausible that USP43 interacts with different 14-3-3 proteins depending on the context. In addition, as 14-3-3 proteins interact with phosphorylated ligands (Fu *et al*, 2000), phosphorylation is likely to provide the recruitment signal, in keeping with our observation that Ser phosphorylated USP43 bound 14-3-3 proteins.

The role of 14-3-3 proteins in HIF signalling has not been well explored. 14-3-3ζ has been implicated in influencing HIF-1α expression (Tang *et al*, 2015; Xue *et al*, 2016), but we did not observe any changes in HIF-1α levels. Instead, the 14-3-3ζ/ε isoforms control the nuclear pool of the HIF-1 complex. 14-3-3 proteins have mostly been associated with enhancing nuclear export (Brunet *et al*., 2002). However, the rapid changes in HIF-1α levels on chromatin suggest that the USP43/14-3-3 interaction with HIF may help with the initial nuclear localisation of the transcription factor. While it will be of interest in future studies to determine how 14-3-3 proteins control nuclear import or delayed export, our work supports phosphorylation as the signal for the association of USP43 with 14-3-3 proteins and HIF-1. Thus, further levels of post-translational regulation can influence HIF-1 signalling, beyond the inhibition of prolyl-hydroxylation and VHL-mediated ubiquitination.

The finding that USP43 was a HIF target gene supported the involvement of this DUB in regulation of a HIF-1 response. However, why USP43 should just regulate HIF-1 signalling, and of these, only a subset of HIF-1 target genes, is less clear. The association of USP43 with HIF-1α and 14-3-3 proteins may partly explain the specificity. It is also possible that USP43 may serve dual functions, in both helping maintain the nuclear pool of HIF-1α via 14-3-3 recruitment, alongside a potential catalytic role of USP43 on chromatin. Recruiting a DUB via a transcription factor may facilitate local chromatin modifications via 14-3-3 (Winter *et al*, 2008) or histone deubiquitination (Peña-Llopis *et al*, 2012). While we did not observe any global change in H2BK120 ubiquitination following USP43 depletion, as previously reported (He *et al*., 2018), it is possible that USP43 may remove other histone ubiquitin marks.

Deubiquitination as a mechanism to reverse the action of VHL has gained considerable focus (Gao *et al*., 2021; Li *et al*., 2005; Mennerich *et al*, 2019; Troilo *et al*., 2014; Wu *et al*., 2016), but the relative importance of individual DUBs in this context is unclear. A single DUB that significantly counteracted VHL-mediated ubiquitination was not identified in our suppressor screen, as OTUD5 loss only had a very marginal effect on the HIF^ODD^GFP reporter in 21 % oxygen. It is possible that there is redundancy or cell type specificity within DUBs reversing VHL-mediated ubiquitination, and their detection may below the stringent statistical cut off used in our screen, potentially explaining why we did not detect DUB that have been previously implicated. DUBs that counteract VHL may also only be uncovered at lower oxygen concentrations. Alternatively, counteracting VHL-mediated ubiquitination may not be required in most contexts, as the oxygen-sensitivity of prolyl hydroxylation provides the regulation and rate-control within the pathway.

The involvement of USP43 in the HIF-1 hypoxia response highlights that coordinated activation HIF signalling involves multiple layers of regulation to refine the response. While prolyl hydroxylation and VHL-mediated ubiquitination are the essential core regulators of the HIF pathway, specific transcriptional responses are mediated downstream, either through the recruitment of co-activators (Batie *et al*, 2019; Batie *et al*, 2022; Chakraborty *et al*, 2019; Galbraith *et al*, 2013; Liu *et al*, 2022; Ortmann *et al*., 2021), the local chromatin environment (Batie *et al*., 2022), or through HIF isoform interactions (Smythies *et al*, 2019). The association of USP43 with 14-3-3 to facilitate a HIF-1 specific response provides a further layer of complexity in how HIF signalling is controls and helps explain how differential transcription programmes are initiated.

## Structured Methods

## Reagents and Tools Table

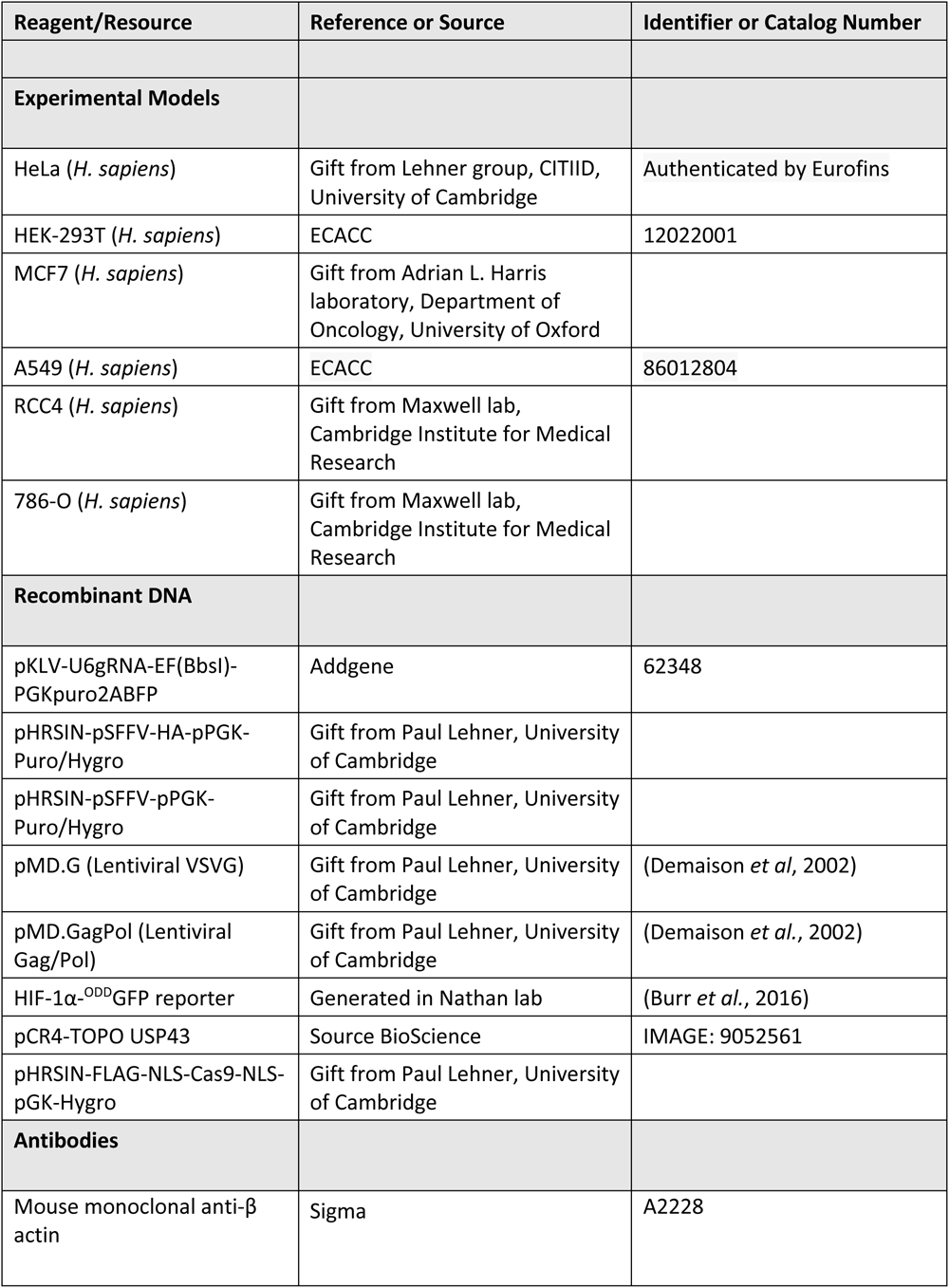
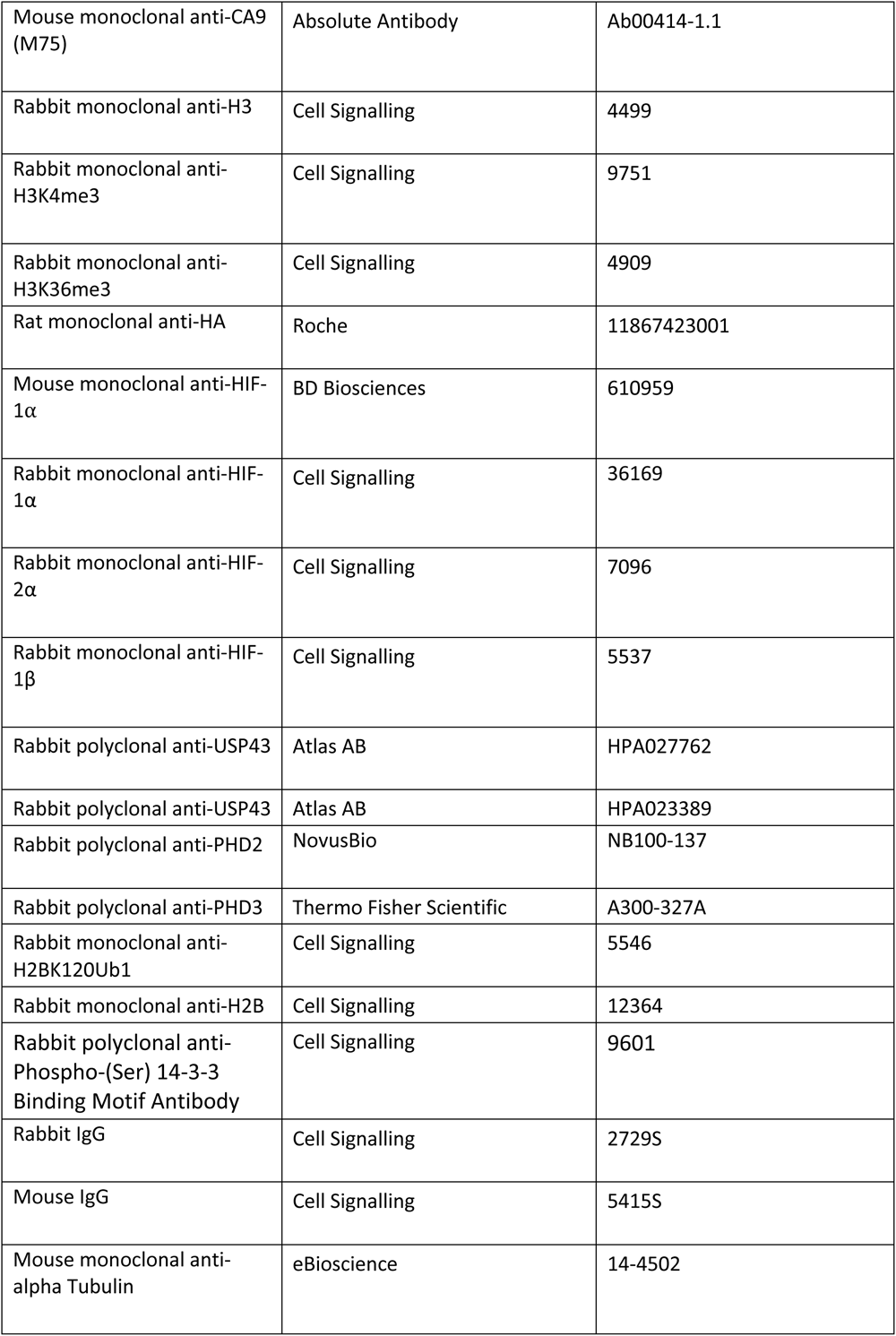
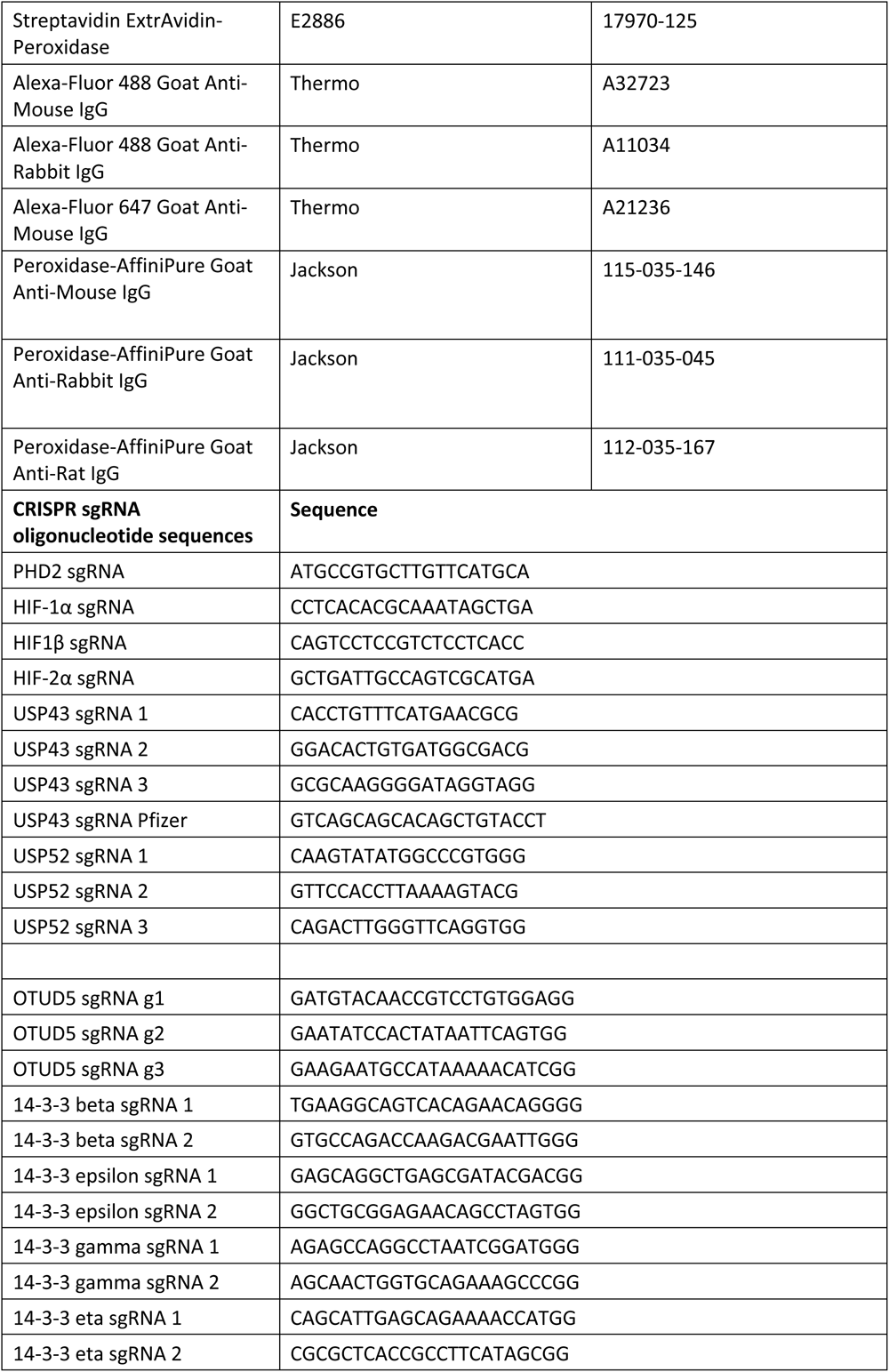
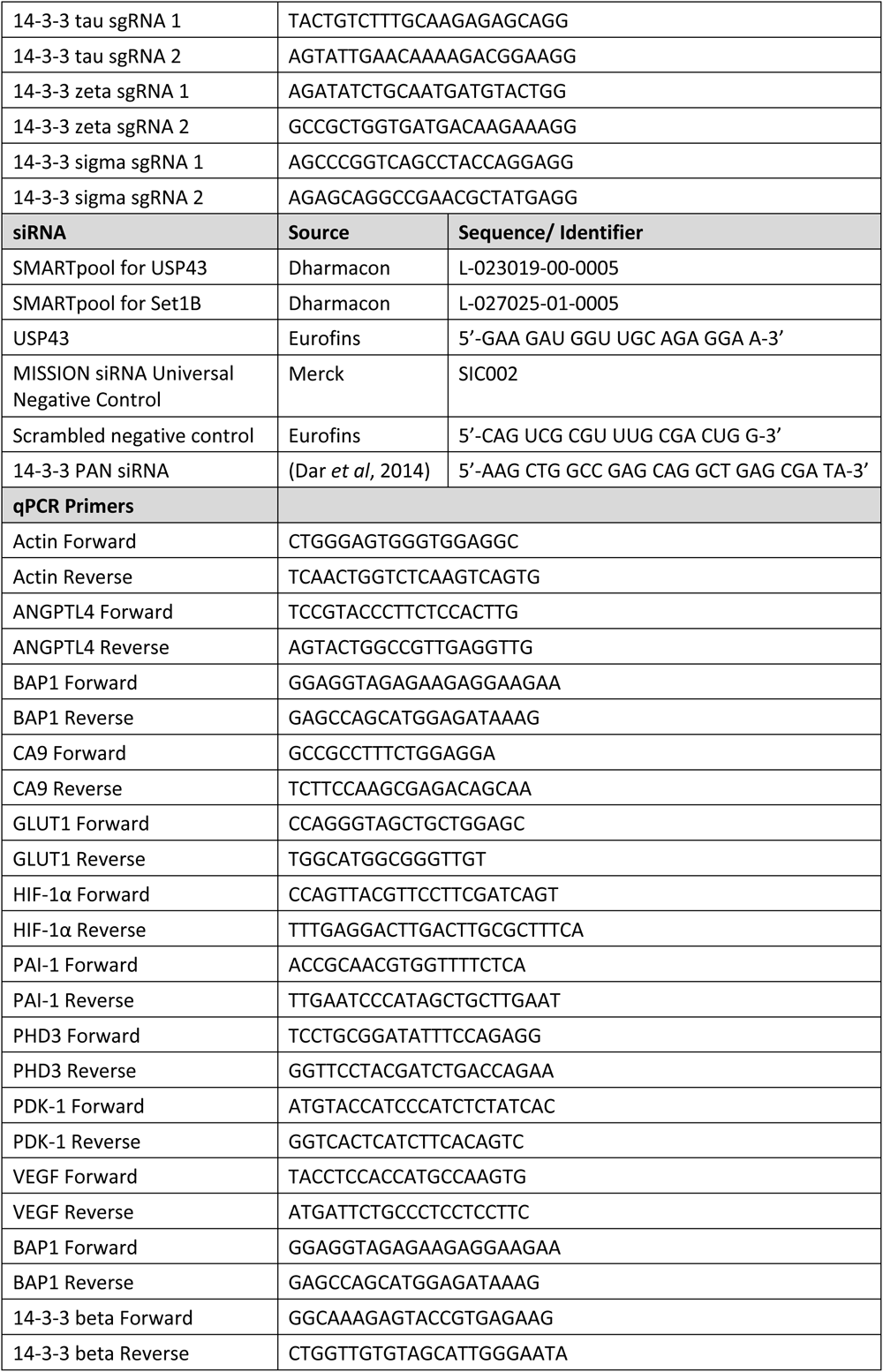
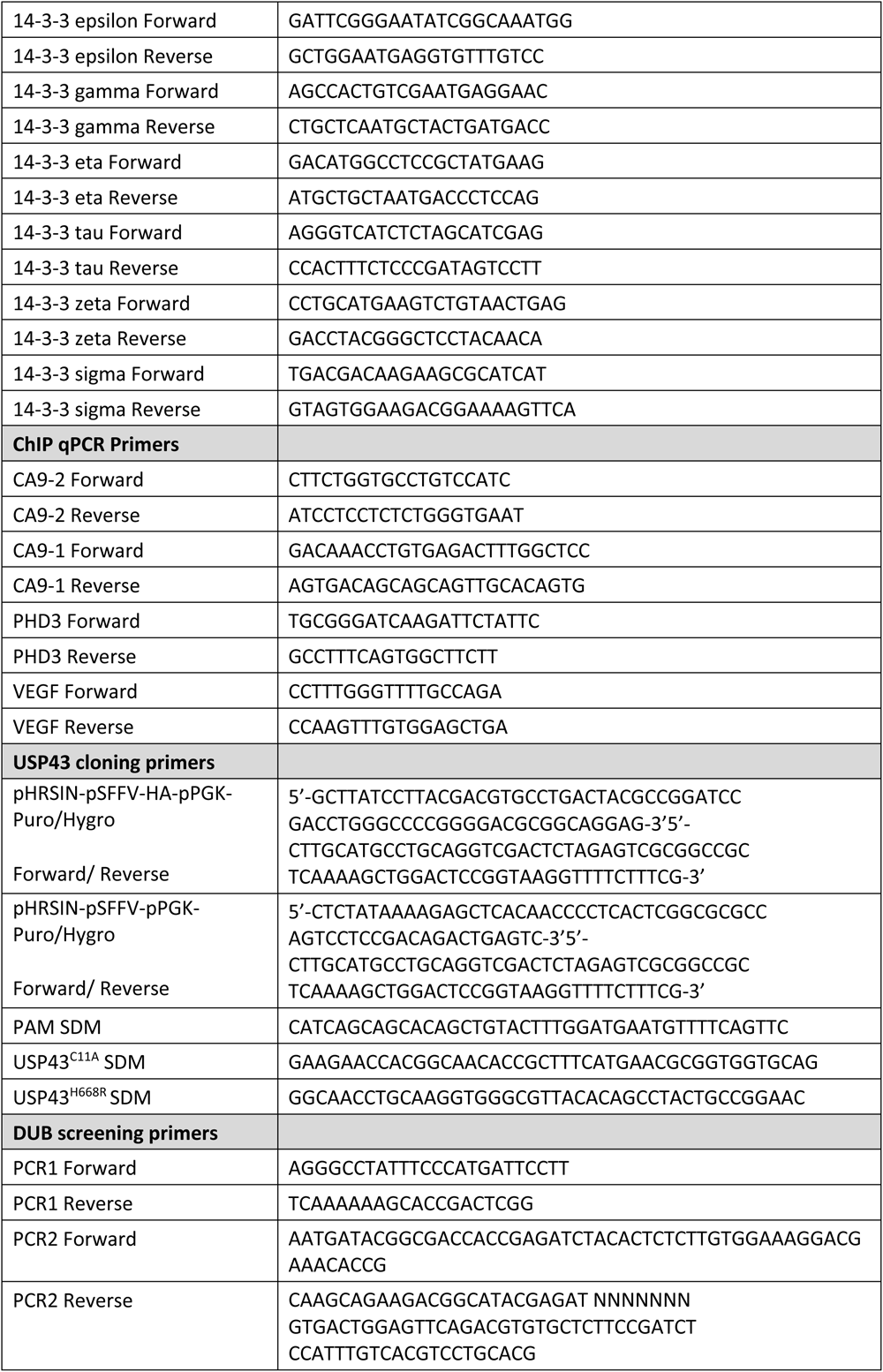
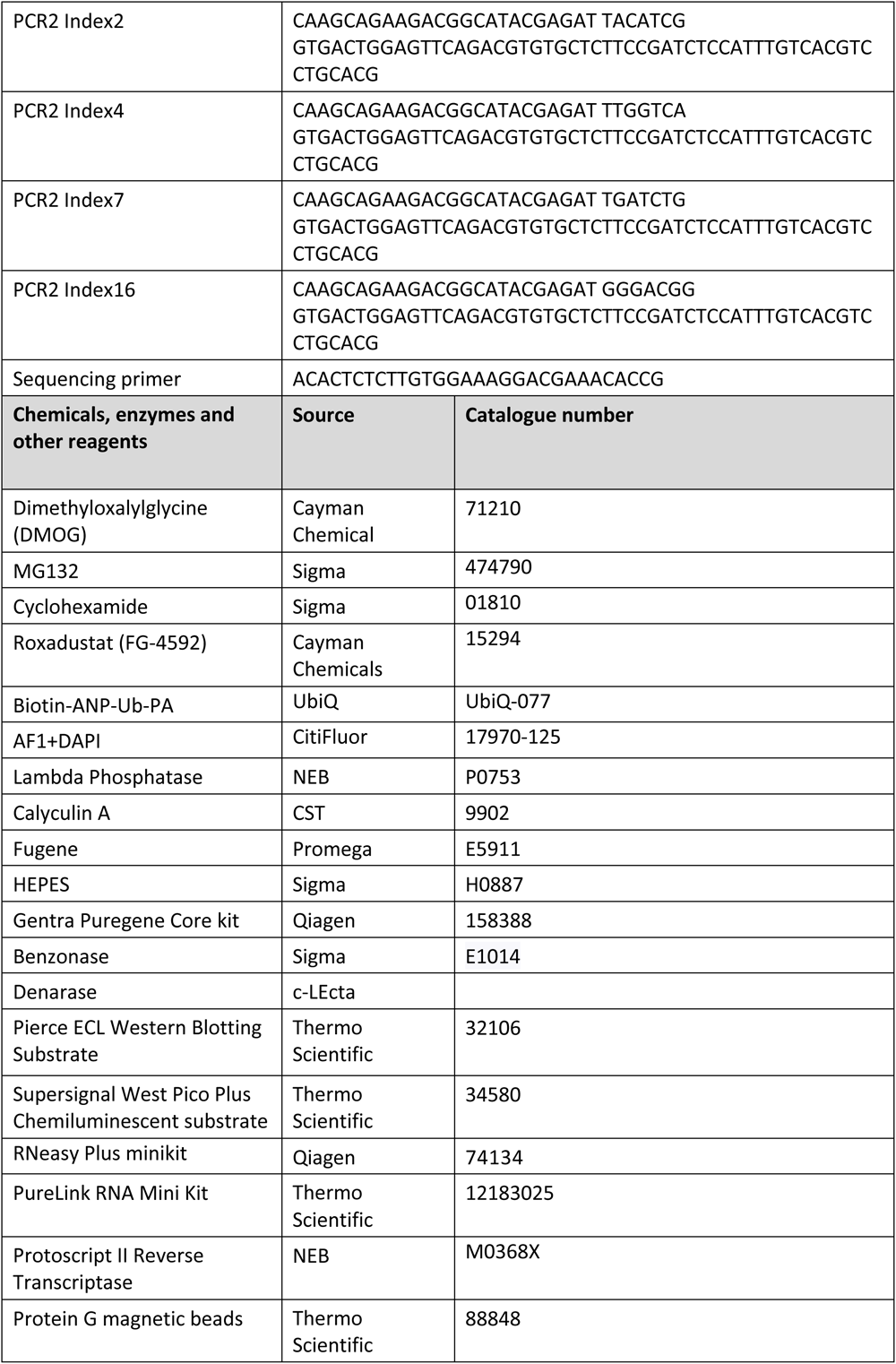
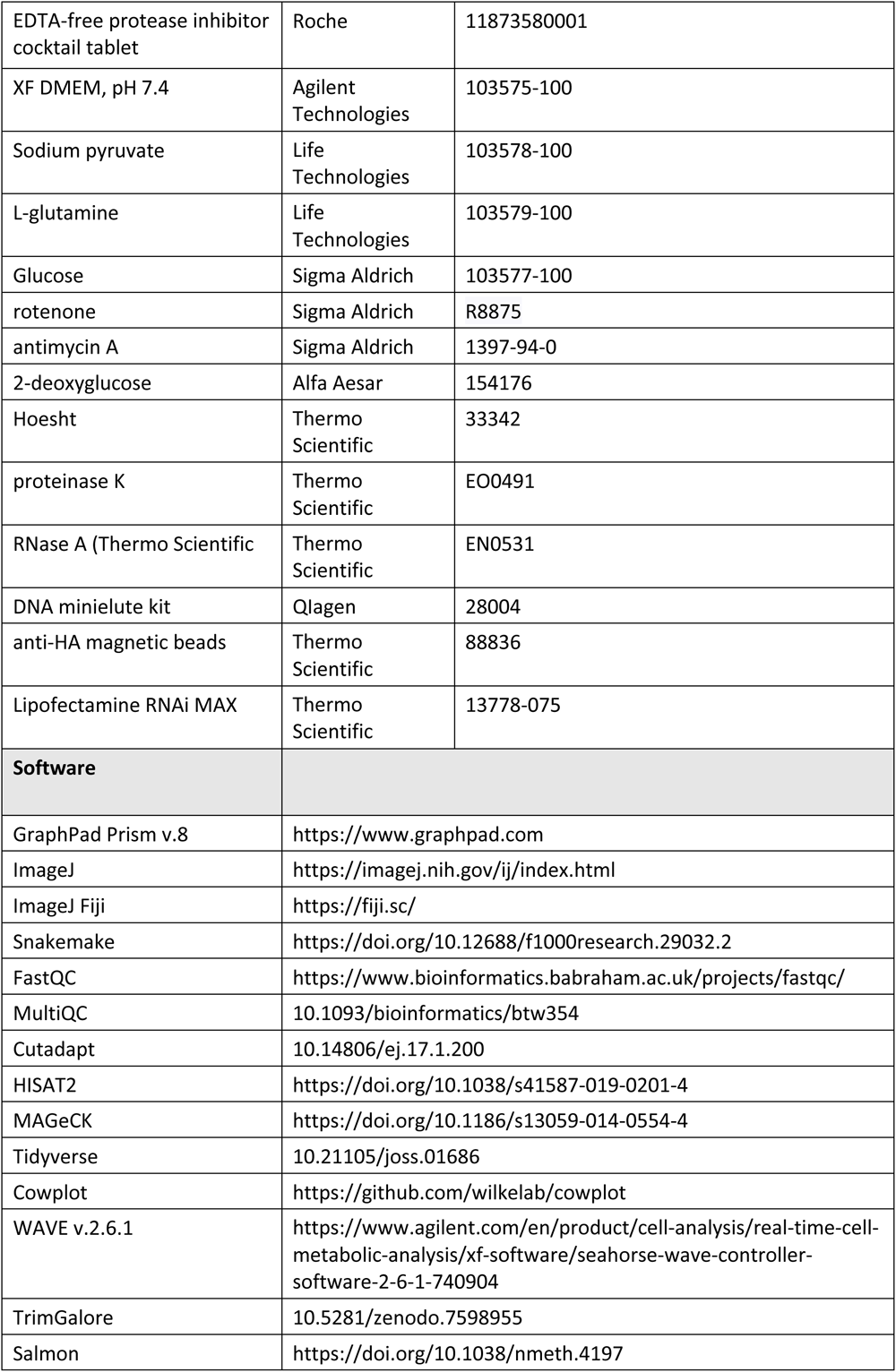
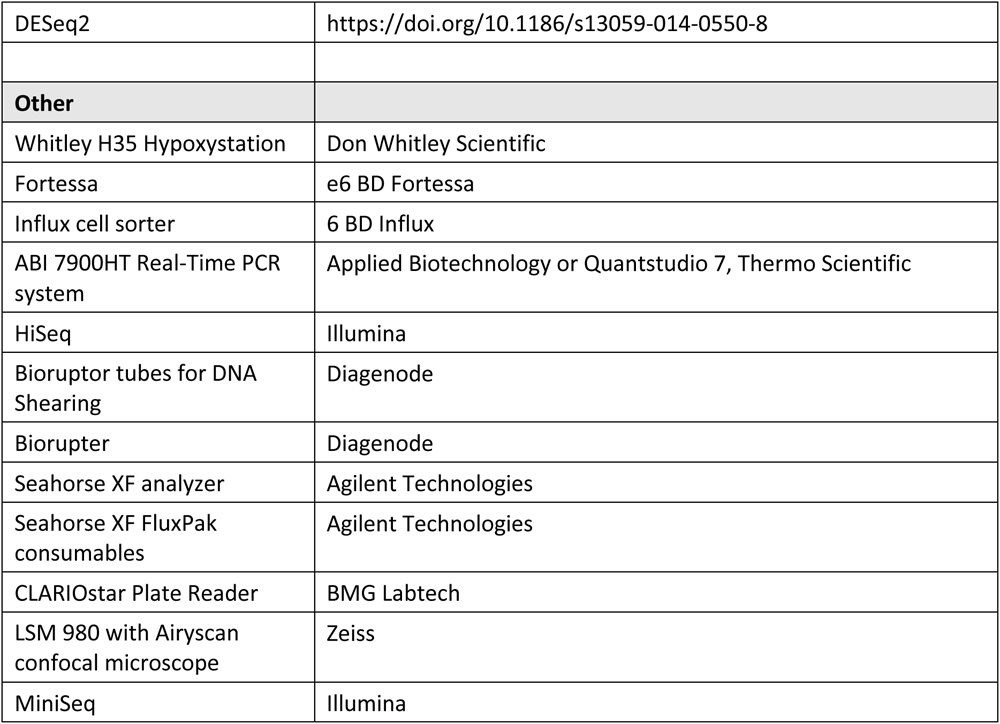

## Methods and Protocols

### Cell culture and reagents

HeLa, HEK293T, MCF7, and A549 cells were maintained in DMEM (Sigma Aldrich) and supplemented with 10% FCS; RCC4 and 786-0 cells were maintained in RPMI-1640 (Sigma) supplemented with 10% FCS. Hypoxic cell culture was performed in a Whitley H35 Hypoxystation (Don Whitley Scientific) at 37 °C, 5% CO_2_, 1 % O_2_ and 94% N_2_. Cells were confirmed mycoplasma negative (Lonza, MycoAltert), and authenticated by short tandem repeat profiling (Eurofins Genomics).

### Plasmids

USP43 constructs were generated from pCR4-TOPO USP43 (Source BioScience, IMAGE:9052561) and cloned into the pHRSIN-pSFFV backbone with puromycin and hygromycin resistance using NEBuilder HiFi (NEB).

### Lentiviral production and transduction

Lentivirus was produced by transfection of HEK293T cells with Fugene (Promega) at 70–80% confluency in six-well plates, with the appropriate pHRSIN vector and the packaging vectors pCMVR8.91 (gag/pol) and pMD.G (VSVG). Viral supernatant was harvested at 48-72 h, filtered (0.22-μm filter), and stored at −80 °C. For transduction, cells were seeded on 24-well plates in 500 µl medium, 500 µl viral supernatant added. Antibiotic selection was applied from 48 h.

### Flow cytometry

A total of 2 × 10^5^ cells per sample were washed in ice-cold PBS in 5-ml FACS tubes and resuspended in 500 µl PBS/formaldehyde before analysis on a BD Fortessa (GFP). For cell-surface staining, cells were washed in ice-cold PBS, incubated at 4 °C for 30 min with the primary antibody, washed with PBS and incubated with the appropriate secondary antibody at 4 °C for 30 min.

### DUB library CRISPR–Cas9 forward genetic screen

The DUB sgRNA library was generated as a subpool of our ‘ubiquitome library’ (Menzies *et al*., 2018). Clonal HeLa HRE-^ODD^GFP cells were transduced with *Streptococcus pyrogenes* Cas9 (pHRSIN-FLAG-NLS-Cas9-NLS-pGK-Hygro) and selected for Cas9 expression using hygromycin. A total of 6 × 10^6^ HeLa HRE-^ODD^GFP cells were transduced with pooled sgRNA virus (multiplicity of infection of around 0.3), establishing at least 500-fold sgRNA coverage, which was maintained throughout the experiments. After 30 h, cells were treated with puromycin 1 μg ml^−1^ for 5 days. The library was pooled before any selection event. After 8 days, we performed FACS by harvesting and sorting 10^8^ cells after 24 h of 1 % O_2_, washing the cells in PBS and resuspending in PBS with 2% FCS and 10 mM HEPES. Cells were sorted using an Influx cell sorter and kept on ice through this process to maintain stability of the reporter. GFP^Low^ cells were chosen in a gate set at 1 log_10_ unit below the mode of the 1 % O_2_ treated control population. Cells were then harvested for DNA extraction. Genomic DNA was extracted using a Gentra Puregene Core kit. Lentiviral sgRNA inserts were amplified in a two-step PCR (with Illumina adapters added on the second PCR).

A Snakemake pipeline (<u>10.5281/zenodo.10286661</u>): after investigating the quality of the raw data using FastQC and MultiQC, reads were quality trimmed to 20 bp using Cutadapt. Trimmed reads were then aligned to all sgRNA sequences using HISAT2 (> 90% alignment rates for all samples). To determine the changes in normalised read counts between samples, MAGeCK was used. The analysis presented compares DNA extracted following the sort to an unsorted DNA library taken at the same time point (**Supplementary Data 1**). All plots showing MAGeCK output were generated with a custom script using the R packages Tidyverse and Cowplot. All scripts used to analyse data and generate plots are publicly available (<u>10.5281/zenodo.10229339</u>).

### CRISPR–Cas9 targeted deletions

Gene-specific CRISPR sgRNA sequences were taken from the Vienna library (www.vbc-score.org) or for USP43, a gift from Pfizer, with 5ʹCACC and 3ʹCAAA overhangs, respectively. SgRNAs were ligated into the pKLV-U6gRNA(BbsI)-PGKpuro2ABFP vector and lentivirus produced as described. Transduced cells were selected with puromycin, and were generally cultured for 9–10 days before subsequent experiments to allow sufficient times for depletion of the target protein. Mixed KO populations were used (denoted by sgRNA) unless otherwise specified. USP43 null clones were also generated from the sgRNA-targeted populations (denoted by KO). Clones were isolated by serial dilution or FACS. USP43 loss was confirmed by immunoblot.

### Immunoblotting

Cells were lysed in an 1xSDS lysis buffer (1 % SDS, 50 mM Tris pH 6.8, 10 % glycerol, 0.01 % (w/v) bromophenol blue, 0.1 M DTT, and 1:500 Benzonase or Denarase for 10 min before heating at 90 °C for 5 min. Proteins were separated by SDS–PAGE, transferred to PVDF (polyvinylidenedifluoride) membranes, probed with appropriate primary and secondary antibodies and developed using enhanced chemiluminescent or Supersignal West Pico Plus Chemiluminescent substrate.

### Quantitative Real Time PCR (qRT-PCR)

Total RNA was extracted using the RNeasy Plus minikit or PureLink RNA Mini Kit following the manufacturer’s instructions and then reversed transcribed using Protoscript II Reverse Transcriptase (NEB). Template cDNA (20 ng) was amplified using the ABI 7900HT Real-Time PCR system (Applied Biotechnology or Quantstudio 7). Transcript levels of genes were normalized to a reference index of a housekeeping gene (β-actin).

### RNA sequencing analysis

Total RNA was extracted from HeLa cells using the RNeasy Plus minikit. Library preparation and sequencing (HiSeq, Illumina) were undertaken by Genewiz. A Snakemake pipeline was used to analyse the data (<u>10.5281/zenodo.10139567</u>): after investigating the quality of the raw data using FastQC and MultiQC, sequence reads were trimmed to remove adapter sequences and nucleotides with poor quality using TrimGalore. Transcript quantification was performed with Salmon against the Gencode Homo sapiens reference transcriptome/genome (build 44) using a decoy-aware transcriptome index for accurate mapping. A mapping rate of > 90% was observed for all samples. DESeq2 was used for differential transcript analysis. Genes with adjusted P values < 0.01 and log_2_(fold change) > 0.5 were called as differentially expressed genes for each comparison (**Supplementary Data 2**) The volcano plots, PCA plot and sample distance heat map were generated with a custom R script using the Cowplot, DESeq2 and Tidyverse R packages. All scripts used to analyse data and generate plots are publicly available (<u>10.5281/zenodo.10229339</u>).

### Immunoprecipitation

HeLa cells were lysed in 1 % Triton PBS or RIPA (25 mM Tris•HCl pH 7.6, 150 mM NaCl, 1 %, NP-40, 1 % sodium deoxycholate, 0.1 % SDS), with 1× Roche complete EDTA-free protease inhibitor cocktail and 1:1000 Denarase for 30 min at 4 °C. Lysates were centrifuged at 14,000 rpm for 10 min, supernatants collected and precleared with Protein G magnetic beads for 2 hat 4 °C. Supernatants were then incubated with primary antibody either overnight (rotation at 4 °C) or with protein G magnetic beads pre-bound to the primary antibody for an hour in 1x PBS with 0.2% Tween (rotation at 4 °C). Protein G magnetic beads were then added for 2 h, and samples were then washed three times in 1xPBS with 1 % Triton and once in 1xPBS. Bound proteins were eluted in 2× SDS loading buffer, separated by SDS–PAGE and immunoblotted.

For phosphor-Serine (14-3-3 motif) immunoprecipitation and subsequent immunoblotting, 1xPBS was substituted by 1xTBS buffer (50 mM Tris-Cl, pH 8, 150 mM NaCl) in all the wash steps.

### Subcellular fractionation

<u>High salt fractionation.</u> 10 × 10^6^ HeLa cells were washed in PBS, lysed in Buffer A (10 mM HEPES, 1.5 mM MgCl_2_, 10 mM KCl, 0.5 mM DTT and EDTA-free protease inhibitor cocktail tablet, Roche) and incubated with rotation at 4 °C for 10 min. Supernatant containing cytosolic fractions were collected by centrifugation (1,400g for 4 min at 4 °C). The pellets were again washed with Buffer A and collected by centrifugation (1,400g for 4 min at 4 °C) to reduce cytosolic fractions leakage to other fractions. The nuclear pellet was resuspended in Buffer B (20 mM HEPES, 1.5 mM MgCl_2_, 300 mM NaCl, 0.5 mM DTT, 25% glycerol, 0.2 mM EDTA and EDTA-free protease inhibitor cocktail tablet) for 10 min on ice to separate nucleoplasmic and chromatin fractions. Samples were centrifuged at 1,700g for 4 min at 4 °C, separating the soluble nucleoplasm from the insoluble chromatin fraction. The chromatin fraction was solubilized in 2× SDS loading buffer containing 1:500 Benzonase or 1:200 Denarase.

### Chromatin fractionation

A cross-liking chromatin enrichment approach was used (Kustatscher *et al*., 2014) with some modifications. 20 × 10^6^ HeLa cells were washed and scraped in cold PBS. Cells were pelleted by centrifugation at 1,800g for 10 min at 4 °C. The pellets were resuspended in 1ml of cold lysis buffer (15mM TRIS pH 7.5, 60mM KCl, 15mM NaCl, 1mM CaCl_2_, 250mM Sucrose, 0.3% NP-40, Protease Inhibitor Cocktail) and incubated for 30 min at 4 °C with rotation. The cytoplasmic extract was separated by centrifugation at 600g for 5 min at 4 °C and lysed in 2xSDS for immunoblotting. The nuclear fraction was washed twice with 1ml of wash buffer (15mM TRIS pH 7.5, 60mM KCl, 15mM NaCl, 1mM CaCl_2_, 250mM Sucrose) spinning at 600g, 5mins, 4 °C. The pellet was fixed in 500 µL 1 % Formaldehyde in PBS and incubated at 37 °C for 10 minutes, then neutralised with 500 µL 500 mM glycine in PBS at RT for 5 min (600g, 5mins). The RNA was removed from the fraction by incubating with 200 µg/mL RNaseA in 500 µL of wash buffer for 15 min at 37 °C in low adhesion 2 ml Eppendorf tube (600g, 5mins). The pelleted fraction was resuspended in 500 µL of SDS buffer (5 % SDS in 25mM TRIS pH 7.5) and 1.5 ml of urea buffer (8 M urea in 25mM TRIS pH 7.5), mixed by inverting and centrifuged at 16,000g for 30 min at RT. The SDS and urea wash was repeated twice. The translucent pellet was washed in 1 ml of SDS buffer (16,000g for 5 min). The chromatin fraction in 100 µL of SDS buffer was sonicated using Bioruptor 0.5 ml Microtubes for DNA Shearing (10 times, 30 s on/off in a Biorupter). Crosslinkeds were reversed by heating at 95 °C for 45 min, and the protein samples dissolved in 100 µL of 2xSDS lysis buffer with 1:200 Denarase for immunoblotting.

### Bioenergetic analyses

Extracellular acidification rates and oxygen consumption rates were measured as indicators of glycolysis and oxidative phosphorylation respectively, using a Seahorse XF analyzer. We performed a glycolytic rate assay according to the manufacturer’s instructions using Seahorse XF FluxPak consumables (Agilent Technologies). Briefly, A549 cells were depleted of HIF-1β or USP43 by sgRNA. Cells were plated in FluxPak 96-well plates in DMEM plus 10% FBS at a cell density of 2 × 10^4^ (A549) with or without 100 µM Roxadustat and incubated at 37 °C for 24 h. The medium was replaced with XF DMEM, pH 7.4 (Agilent Technologies) supplemented with 1 mM sodium pyruvate, 2 mM L-glutamine, 10 mM glucose and incubated at 37 °C in a non-CO_2_ incubator for 1 h before replacing with the same medium before measuring in the analyser. Program settings: mix 3 min, measure 3 min ×3; inject 0.5 µM rotenone and 0.5 µM antimycin A (; mix 3 min measure 3 min ×3; inject 50 mM 2-deoxyglucose; mix 3 min, measure 3 min ×5. Medium was then removed, and cells quantified using the 100 µl 2.5 µM Hoechst in PBS and CLARIOstar Plate Reader at 355-20/455-30 nm. Results were normalized and analysed using WAVE v.2.6.1 and data exported to the Agilent Seahorse XF Glycolytic Rate Assay Report Generator. Graphs were prepared using GraphPad Prism v.8.

### Chromatin Immunoprecipitation PCR (ChIP-PCR)

HeLa cells were grown on 15-cm dishes to 15 × 10^6^ density, and then treated with 1 % formaldehyde for 10 min to crosslink proteins to chromatin. The reaction was quenched with glycine (0.125 M, 10 min at room temperature). Cells were washed in ice-cold PBS twice, scraped in tubes, and centrifuged at 800 rpm for 10 min before lysis in 500 µl of ChIP lysis buffer (50 mM Tris-HCl (pH 8.1), 1 % SDS, 10 mM EDTA, Complete Mini EDTA-free protease inhibitor). Samples were incubated on ice for 10 min and diluted 1:1 with ChIP dilution buffer (20 mM Tris-HCl (pH 8.1), 1 % (v/v) Triton X-100, 2 mM EDTA and 150 mM NaCl). Samples were then sonicated for 20 cycles of 30s on and 30s off in a Biorupter, followed by centrifugation for 10 min at 13,000 rpm at 4 °C. Supernatants were collected and 20 µl stored at −20 °C as the input sample. A 200 µl aliquot of the remaining sample was diluted with ChIP dilution buffer to 1 ml, and precleared using 25 µl Protein G magnetic beads (4 °C, 2 h, rotating); 1 ml of sample was immunoprecipitated with the appropriate primary antibody (4 °C, overnight, rotating). Protein G magnetic beads (25 μl) were added and incubated for a further 2 h at 4 °C. The beads were washed sequentially for 5 min each with wash buffer 1 (20 mM Tris-HCl (pH 8.1), 0.1 % (w/v) SDS, 1 % (v/v) Triton X-100, 2 mM EDTA, 150 mM NaCl), wash buffer 2 (Wash Buffer 1 with 500 mM NaCl), wash buffer 3 (10 mM Tris-HCl (pH 8.1), 0.25 M LiCl, 7 1 % (v/v) NP-40, 1 % (w/v) Na-deoxycholate and 1 mM EDTA), and twice with TE buffer (10 mM Tris-HCl (pH 8.0) and 1 mM EDTA). Bound complexes were eluted with 120 μl elution buffer (1 % (w/v) SDS and 0.1 M Na-bicarbonate), and crosslinking reversed by addition of 0.2 M NaCl, and incubation at 65 °C overnight with agitation (300 rpm). Protein was digested with 20 μg proteinase K for 4 h at 45 °C. RNase A was added for 30 min at 37 °C and DNA purified using the DNA minielute kit. DNA underwent qPCR analysis, and results expressed relative to input material.

### siRNA-mediated depletion

HeLa or A549 cells were transfected with siRNA SMARTpools for USP43 (Dharmacon, L-023019-00-0005), USP43 siRNA (Eurofins) (5’-GAA GAU GGU UGC AGA GGA A-3’), MISSION siRNA Universal Negative Control (SIC002, Merck), Scrambled negative control (Eurofins): (5’-CAG UCG CGU UUG CGA CUG G-3’), or 14-3-3 PAN siRNA (5’-AAG CTG GCC GAG CAG GCT GAG CGA TA-3’) (Dar *et al*., 2014) using Lipofectamine RNAi MAX. Cells were harvested after 48-72 h for further analysis by flow cytometry, qPCR or immunoblot.

### USP43 transfection or transduction

pHRSIN-pSFFV-pPGK-Puro/Hygro USP43 expressing vector was transfected to 5x10^5^ cells in 6 well plates using Fugene. Alternatively, lentivirus was harvested from the transfected cells, as described above. A calcium phosphate (CaPO_4_) transfection protocol was used to overexpress HA-USP43 constructs for DUB activity assay. HEK293T cells at 40 % confluency in 10 ml of DMEM were transfected with 4 µg of plasmid. CaCl_2_/HBS/DNA precipitate was prepared by mixing 438 µl of H_2_O with 2M CaCl_2_ and DNA in one tube and then adding it dropwise to 500 µl of 2xHBS (0.27M NaCl, 1.5 mM Na_2_HPO_4_·7H_2_O, 0.055M HEPES, pH 7). The mix was transferred to cells by distributing dropwise all around the dish and gently swirling the dish around. The media was changed the following day and cells were harvested 48 h after transfection.

### DUB activity probe assay

HA-USP43, HA-USP43^C110A^, or HA^C110A+H667R^ were overexpressed in HEK293T cells using the calcium phosphate transfection protocol. Cells were scraped in 500 µl of 1 % Triton in 1xPBS, and centrifuged for 10 min at 14,000 g at 4 °C. Input samples were taken at this point. The remaining samples were pre-cleared with 50 µl sepharose beads for 2 hours at 4 °C, and incubated with 25 µl of anti-HA magnetic beads (at 4 °C overnight. The beads were washed 3 times in 1 % Triton in 1xPBS and once in 1xPBS, resuspended in 100 µl of 1xPBS with 5 µl of DUB catalytic activity-based probe Biotin-ANP-Ub-PA (1 µM final concentration) and incubated at 37 °C for 5 min. The reactions were washed 3 times in 1x PBS, lysed with 50 µl 2 x SDS and analysed by immunoblotting.

### Cycloheximide chase assays

HeLa USP43 KO or overexpressing cells were seeded at 3x10^5^ cells in 6 well plates the night before. The cells were pre-incubated in 1 % oxygen for 4 h and then treated with 100 µg/ml of cycloheximide for 0, 20. 40, and 60 min and analysed by immunoblotting for HIF-1α degradation.

### Phosphorylation-dependent electrophoretic mobility shift (PDEMS) assay

1 x 10^6^ of Hela cells were washed in 1 x TBS buffer (50 mM Tris-Cl, pH 8, 150 mM NaCl), scraped and lysed in 80 µl of RIPA. The lysate was incubated with 1 µl (400 units) of Lambda protein phosphatase (NEB, P0753) according to manufacturer’s instructions. The reaction was stopped with 0.2 µl of 0.5 M EDTA, and samples were prepared for immunoblotting by adding 20 µl 6 x SDS loading dye and boiling.

### Immunofluorescence

HeLa cells were seeded on FCS precoated coverslips in 24 well plates at 5x10^4^ density. The following day, cells were incubated in 21 or 1 % oxygen for 6 h, washed twice in 1xDPBS and fixed with 4 % Paraformaldehyde for 15 min. Cells were washed twice in 1xDPBS, permeabilised with 0.1 % Triton for 10 min, washed again, and blocked in 4 % FBS in 1xDPBS. The coverslips were stained with 1:100 of anti-USP43, or anti-HIF-1α primary antibodies at 4 °C overnight, then washed 5 times with 1xDPBS and counterstained with 1:400 anti-mouse 647 nm, or anti-rabbit 488 nm Alexa Fluor™ secondary antibodies for 1.5 h at room temperature. The cells were washed 5 times in 1x DPBS and mounted in glycerol based mountant with DAPI. The cells were imaged using LSM 980 with Airyscan confocal microscope and analysed blinded with Fiji (ImageJ-win64).

### Statistical analyses

Quantification and data analysis of experiments are expressed as mean ± sd and *P* values were calculated using analysis of variance (ANOVA) or two-tailed Student’s *t*-test for pairwise comparisons using Graphpad Prism v.8. Qualitative experiments were repeated independently to confirm accuracy. Statistical analyses for the CRISPR screens, RNA-seq and ChIP-seq are described in the relevant sections.

## Acknowledgements

We thank all members of the Nathan lab for their helpful comments on the work and manuscript. This work was supported by a Pfizer ITEN award to JAN (Pfizer Inc), a Wellcome Senior Clinical Research Fellowship to JAN (215477/Z/19/Z), and a Lister Institute Research Fellowship to JAN. This work was also supported by the NIHR BRC.

## Author Contributions

Conceptualization, TP and JAN; Methodology, TP, NW, RVS and JAN; Investigation, TP, NW, RVS and JAN; Writing – original draft, TP and JAN; Writing – reviewing and editing, all authors; Funding acquisition, JAN; Resources, JAN; Supervision, JAN.

## Conflict of Interest

JAN received a Pfizer ITEN discovery grant to fund this work. Other authors declare no competing interests.

## Notes

### Competing Interest Statement

JAN is receipt of a Pfizer ITEN award that contributed funding to this work.

### Summary of Updates

Supplementary Information has been included.

