## Supplementary Information for "Deubiquitinating enzyme mutagenesis screens identify a USP43 driven HIF-1 transcriptional response"

**Fig S1. Validation of DUBs identified as regulators of the HIF response in the DUB screens.**

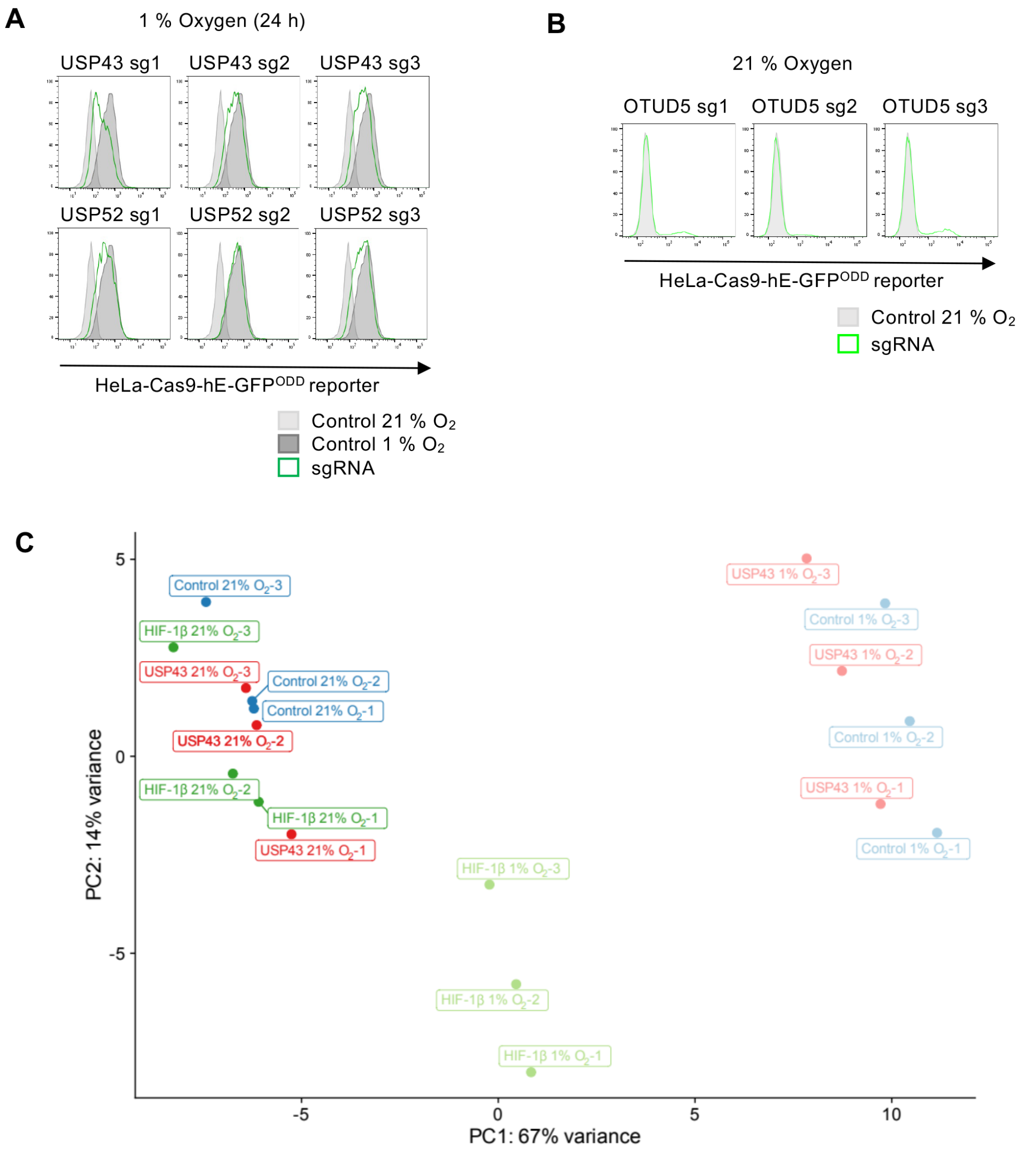


**A)** HeLa HRE-$^{ODD}$GFP reporter cells transduced with 3 different sgRNAs against USP43, or USP52 were generated. Cells were then incubated in 1 % oxygen for 24 h and analysed by flow cytometry. Representative of three biological replicates. **B)** HeLa HRE-$^{ODD}$GFP reporter cells transduced with 3 different sgRNAs against OTUD5 cultured in 21 % oxygen. Representative of three biological replicates **C)** Principal component analysis (PCA) plot of RNA-seq of HeLa control, HIF1β KO, and USP43 KO cells that were treated with 21 % or 1 % oxygen ($O_2$) for 16 h before RNA was extracted and sequenced using Hi-seq. n= 3 biologically independent samples.

**Fig S2. USP43 depletion delays activation of a HIF response.**

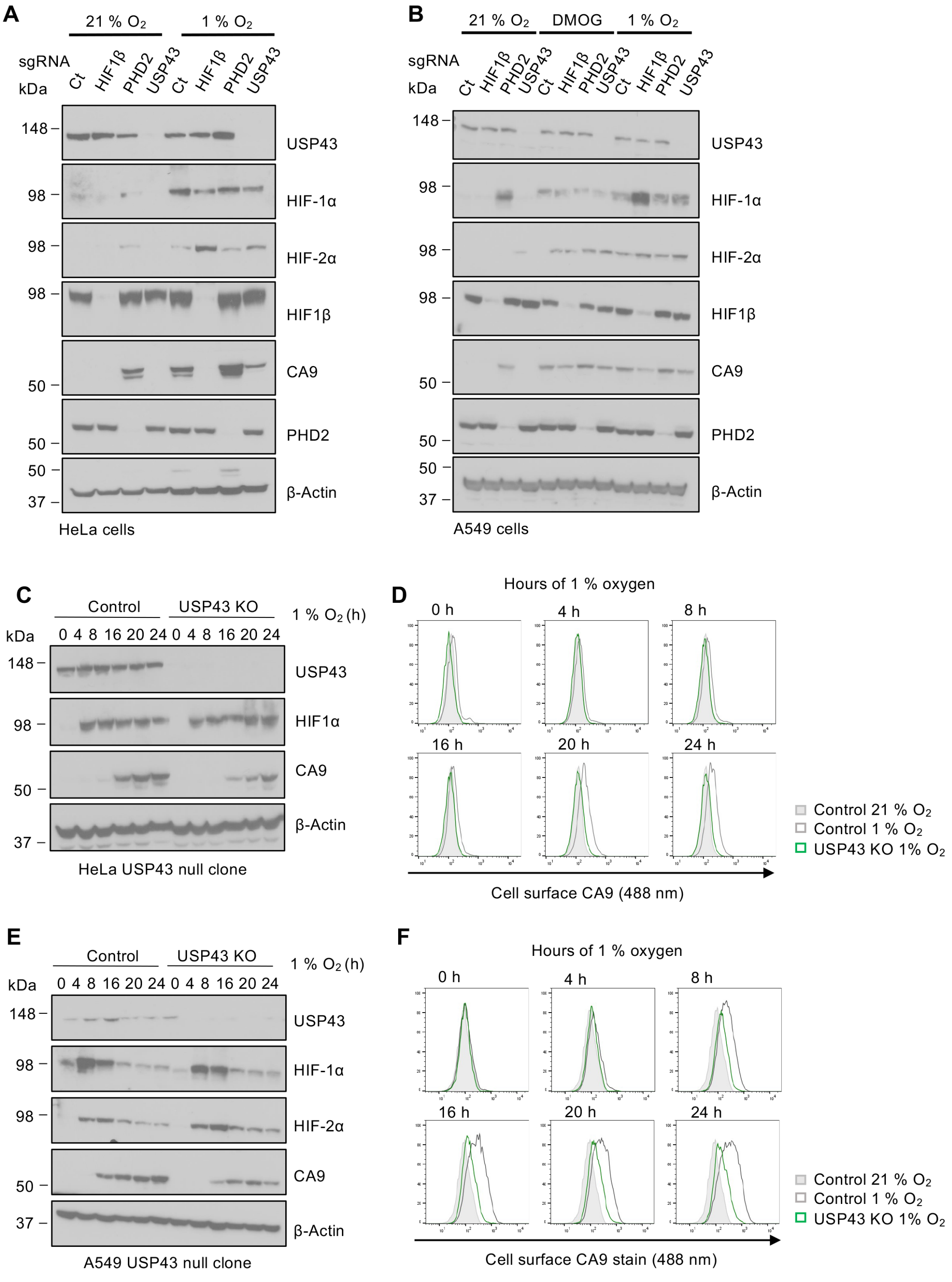


**A)** Control (Ct) or mixed KO populations of HIF1β, PHD2, or USP43 KO HeLa cells were incubated in 21 % or 1 % oxygen and immunoblotted for components of the HIF pathway and the HIF-1 target, CA9. Representative of three biological replicates. **B)** Control or mixed KO populations of HIF1β, PHD2, or USP43 KO A549 cells were incubated in 21 %, 1 % oxygen, or treated with 1 mM DMOG, and immunoblotted for components of the HIF pathway and the HIF-1 target, CA9. Representative of three biological replicates. **C)** HeLa control cells or a HeLa USP43 null clone were incubated in 0-24 h of 1 % oxygen and analysed by immunoblotting. Representative of three biological replicates. **D)** Cell surface

CA9 levels in control or mixed KO populations of USP43 HeLa cells incubated in 0-24 h of 1 % oxygen. Plots are representatives of three biological replicates. **E, F)** As for **(C, D)** but using A549 cells. Representative of three biological replicates.

**Fig S3. USP43 depletion decreases the expression of selected HIF target genes in hypoxia.**

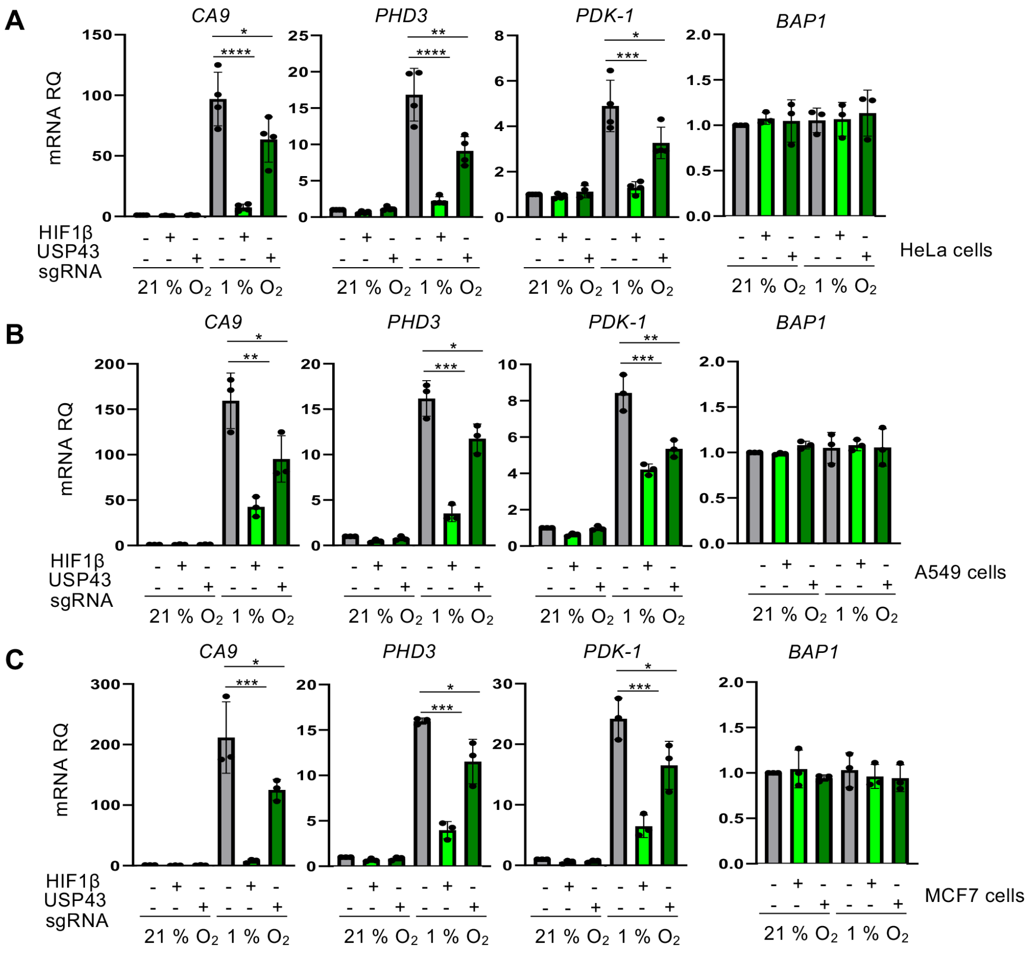


**A-C)** RT-qPCR of *CA9*, *PHD3*, *PDK1*, and *BAP1* in HeLa **(A)**, A549 **(B)** and MCF7 **(C)** cells with or without depletion of HIF1β or USP43. Cells were incubated in 21 % or 1 % oxygen for 16 h prior to lysis. n=4 biologically independent samples, mean ± sd, *P≤ 0.05, **P≤ 0.01, ***P≤ 0.001, ****P≤ 0.0001, two-way ANOVA.

**Fig S4. USP43 is specific for the HIF-1 complex.**

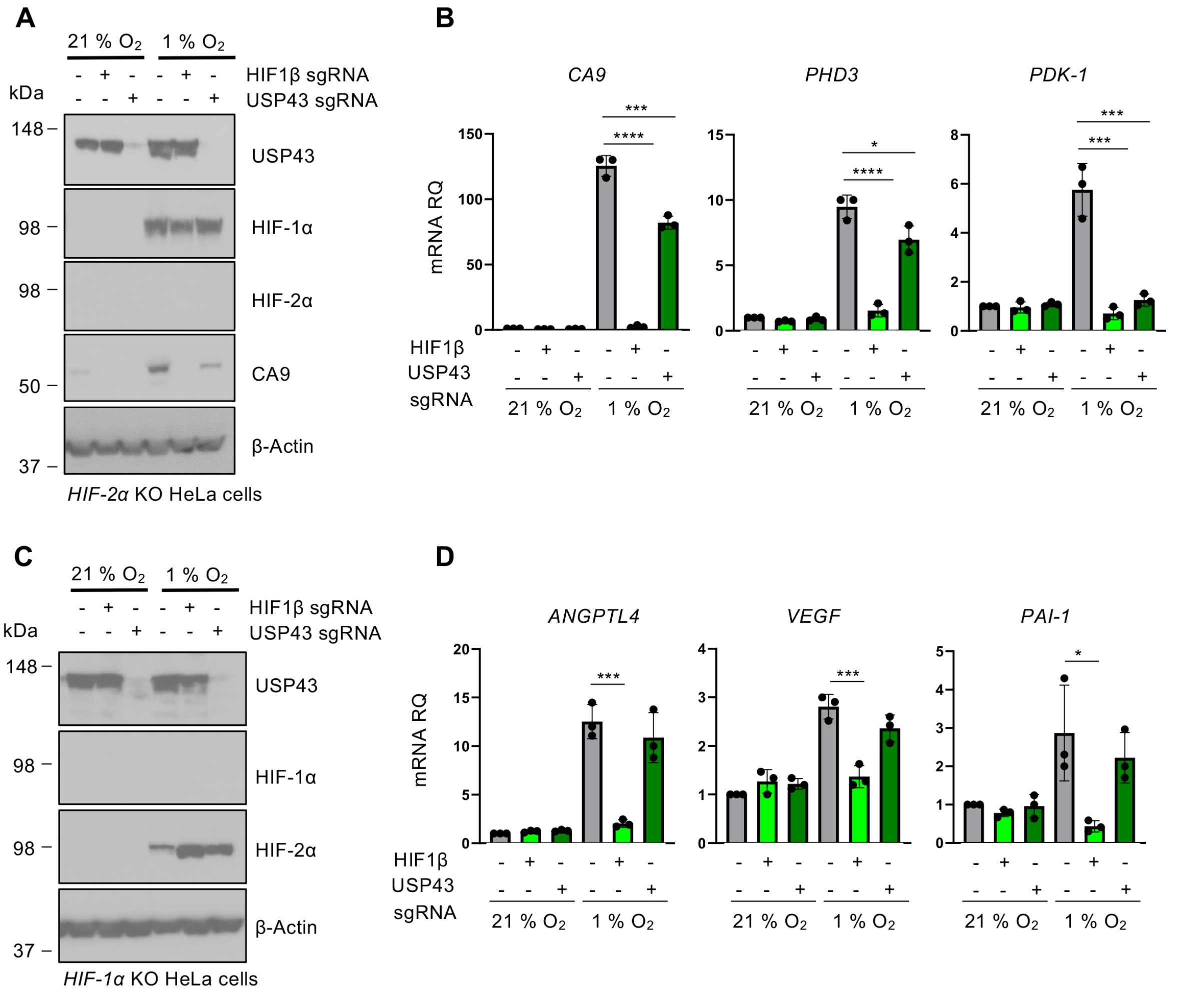


**A, B)** Clonal HIF-2α null HeLa cells were transduced with sgRNA targeting HIF1β or USP43. Cells were incubated in 21 % or 1 % oxygen for 16 h and analysed by immunoblot **(A)** or qPCR for selected HIF target genes **(B)**. n=3 biologically independent samples, mean ± sd, *P≤ 0.05, ***P≤ 0.001, ****P≤ 0.0001, two-way ANOVA. **C, D)** Clonal HIF-1α null HeLa cells were transduced with sgRNA targeting HIF1β or USP43. Cells were incubated in 21 % or 1 % oxygen for 16 h and analysed by immunoblot **(C)** or qPCR for selected HIF target genes **(D)**. n=3 biologically independent samples, mean ± sd, *P≤ 0.05, ***P≤ 0.001, two-way ANOVA.

**Fig S5. Reconstituting USP43 deficiency restores HIF signalling.**

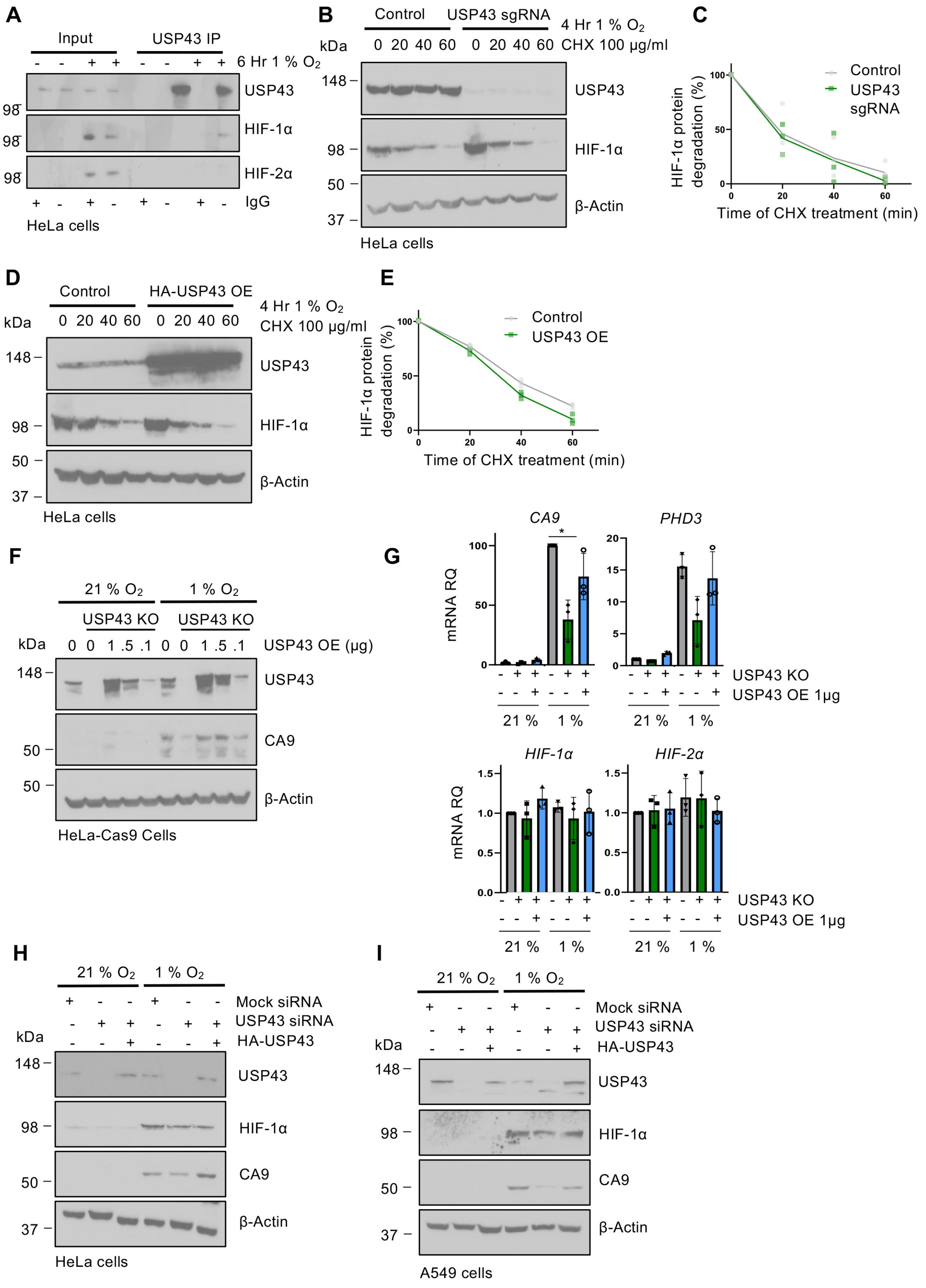


**A)** Endogenous USP43 was immunoprecipitated in HeLa cells grown in 21 % or 1 % oxygen for 6 h. Samples were immunoblotted for HIF-1α and HIF-2α. Representative of 3 biological replicates. **B, C)** Control or mixed population USP43 KO HeLa cells were incubated in 1 % oxygen for 4 h and then treated with cycloheximide (100 µg/ml) in hypoxia for 0-60 min. HIF-1α levels were measured by immunoblot **(B)** and quantified by densitometry using ImageJ **(C)**. **D, E)** n=3 biological replicates.

Control or USP43 overexpressing (OE) HeLa cells were incubated in 1 % oxygen for 4 h and then treated with cycloheximide (100 µg/ml) in hypoxia for 0-60 min. HIF-1$\alpha$ levels were measured by immunoblot **(D)** and quantified by densitometry using ImageJ **(E)**. **F, G)** Control or USP43 clonal KO cells were reconstituted with 0, 1, 0.5, or 0.1 µg by transient transfection of USP43, incubated in 21 % or 1 % oxygen for 16 h, and analysed by immunoblot **(F)** or qPCR **(G)**. Representative of three biological replicates. n=3 biologically independent samples, mean ± sd, *P≤ 0.05, two-way ANOVA. **H, I)** USP43 was depleted in HeLa cells using siRNA and compared to a mock siRNA control. siRNA transfected HeLa **(H)** or A549 **(I)** cells were then transfected with HA-USP43 (1 µg) to reconstitute USP43, and incubated in 21 % or 1 % oxygen for 16 h. Immunoblot representative of three biological replicates.

**Fig S6. USP43 regulates HIF-1α nuclear accumulation.**

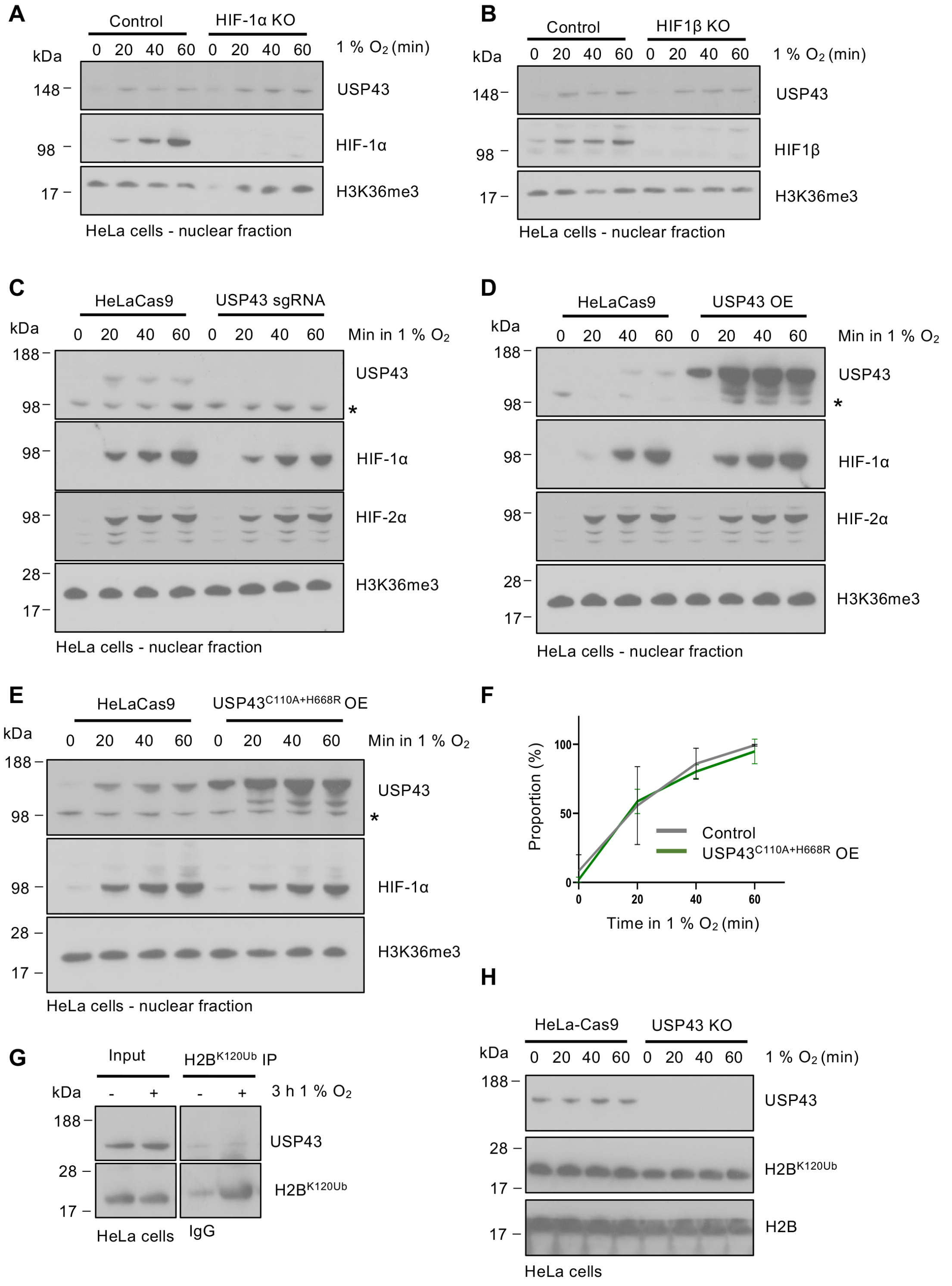


**A)** Immunoblot of the nuclear fraction of HeLa-Cas9 or HIF-1α KO cells incubated in 1 % oxygen for 0-60 min. Samples were immunoblotted for USP43, HIF-1α, and H3K36me3 as a loading control. Representative of 3 biological replicates. **B)** As for **(A)** but in HIF1β KO cells. Representative of 3 biological replicates. **C-F)** Immunoblot of the nuclear fraction of HeLa-Cas9 or mixed population USP43 KO cells **(C)**, USP43 overexpressing (OE) cells **(D)**, or USP43$^{C110A+H668R}$ OE cells **(E)** incubated in 1 %

oxygen for 0-60 min. Representative of 3 biological replicates. Quantification of HIF-1α enrichment within the nuclear fraction in HeLa control or USP43$^{C110A+H668R}$ OE cells **(F)**. HIF-1α levels relative to a stable histone mark (H3K36me3) were measured by immunoblot. n=3 biologically independent samples, mean ± sd, two-way ANOVA. **G)** Endogenous H2B$^{K120Ub}$ was immunoprecipitated in HeLa cells grown in 21 % or 1 % oxygen for 3 h. Samples were immunoblotted for USP43. Representative of 3 biological replicates). **H)** Immunoblot of H2B$^{K120Ub}$ levels in control or USP43 null HeLa cells incubated in 1 % oxygen for 0-60 min. Representative of 3 biological replicates. **represents non-specific band.*

**Fig S7. USP43 associates with 14-3-3 proteins to regulate HIF-1 signalling.**

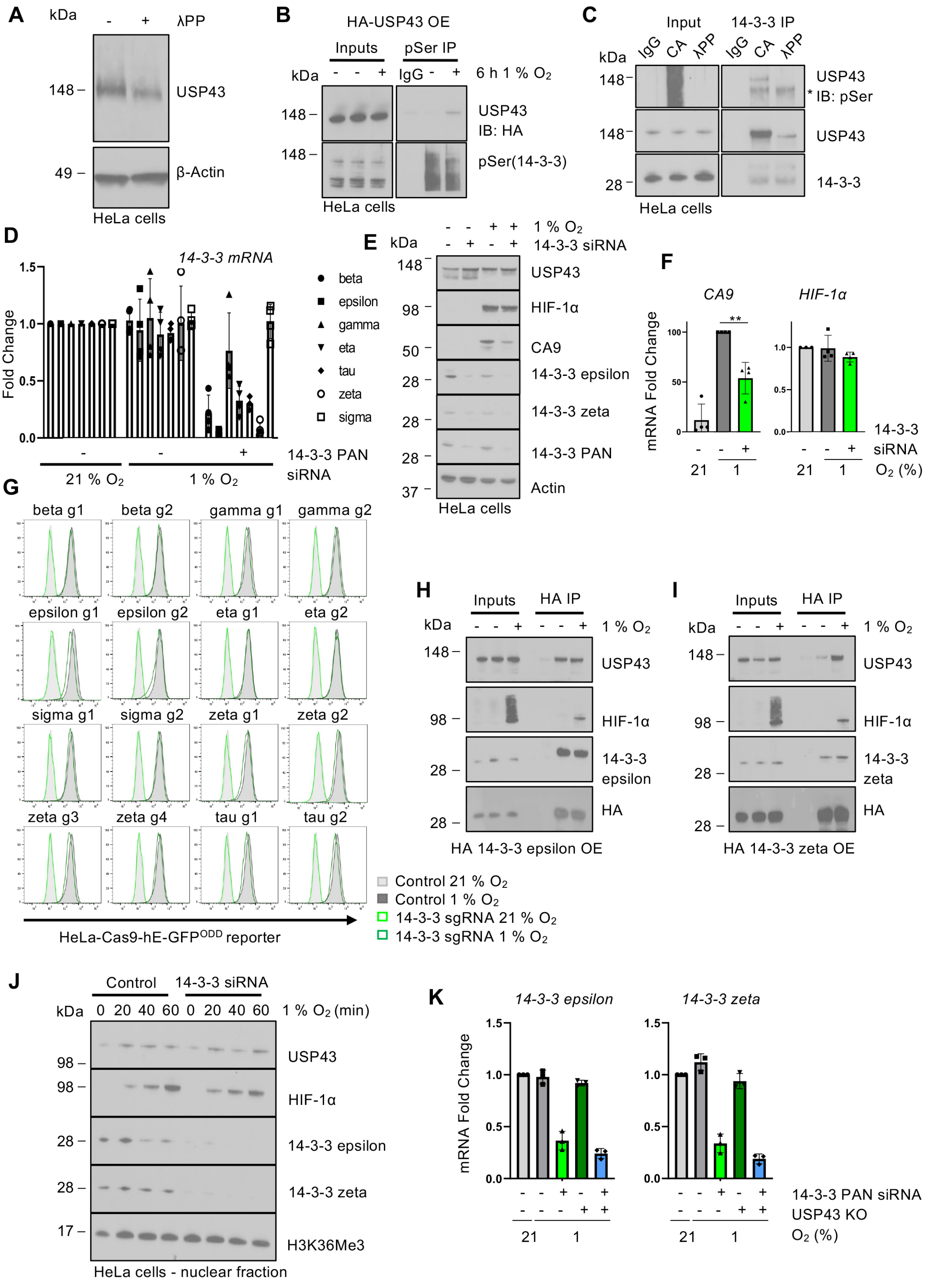


**A)** Immunoblot of the phosphorylation-dependent electrophoretic mobility shift (PDEMS) assay of USP43, with or without Lambda Protein Phosphatase (λPP) treatment. **B)** Endogenous pSer(14-3-3 motif) was immunoprecipitated in HA-USP43 overexpressing HeLa cells incubated in 21 % or 1 % oxygen for 6 h. Representative of 3 biological replicates. **C)** Endogenous 14-3-3 was immunoprecipitated in HeLa cells with or without Calyculin A (CA) treatment (100 nM, 37°C for 30 min) or λPP (400 units, 30°C for 30 min). Samples were immunoblotted for pSer(14-3-3 motif), USP43, and 14-3-3. Representative of 3 biological replicates. **D)** qPCR of HeLa cells transfected with a 14-3-3 pan siRNA, and incubated in 1 % or 21 % oxygen for 16 hr. Expression of the seven *14-3-3* isoforms was analysed using individual primer pairs. n=3 biologically independent samples, mean ± sd. **E)** Immunoblot of control or 14-3-3 siRNA-depleted HeLa cells incubated in 21 % or 1 % oxygen for 16 h. Representative of three biological replicates. **F)** qPCR of *CA9* and *HIF-1α* mRNA expression control or 14-3-3 siRNA-depleted HeLa cells incubated in 21 % or 1 % oxygen for 16 h. n=3 biologically independent samples, mean ± sd, $^{**}P \leq 0.01$, two-way ANOVA. **G)** Mixed KO populations of HeLa HRE-$^{ODD}$GFP reporter cells to each 14-3-3 isoform were generated by lentiviral transduction of sgRNA (2 sgRNAs for each isoforms, 4 sgRNAs for zeta isoform). Cells were incubated in 21 % or 1 % oxygen for 16 h and analysed by flow cytometry. **H, I)** HeLa cells were transfected with HA-14-3-3 epsilon **(H)** or HA-14-3-3 zeta **(I)**, and incubated in 21 % or 1 % oxygen for 6 h. The 14-3-3 isoforms were immunoprecipitated using the HA tag and immunoblotted for USP43. Representative of 3 biological replicates. **J)** HIF-1α enrichment within the nuclear fraction in HeLa control or following 14-3-3 siRNA-mediated depletion. Cells were incubated in 1 % oxygen for 0 to 60 min. **K)** qPCR of *14-3-3 epsilon* and *zeta* expression in control or USP43 null cells, with or without 14-3-3 siRNA-mediated depletion. Cells were incubated in 21 % or 1 % oxygen for 6 hr prior to lysis. n=3, mean ± sd.

### Gating strategy

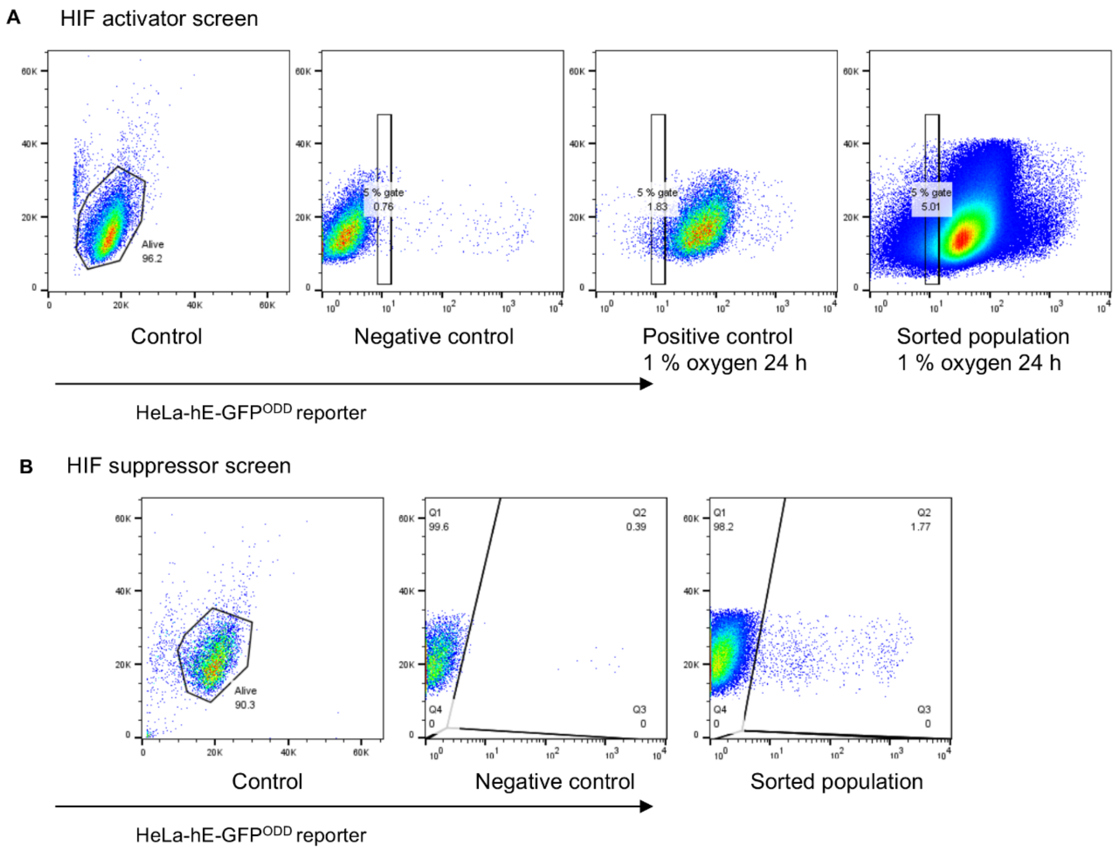


**A, B)** Representative gating strategy for the HeLa-HRE-GFP$^{ODD}$ reporter screens. Gates for sorting the low GFP population in 1% oxygen **(A)** and GFP high population in 21% oxygen **(B)**.
